# Dynamic interactions and intracellular fate of label-free, thin graphene oxide sheets within mammalian cells: role of lateral sheet size

**DOI:** 10.1101/805200

**Authors:** Yingxian Chen, Jack Rivers-Auty, Livia Elena Crică, Katie Barr, Vinicio Rosano, Adrian Esteban Arranz, Thomas Loret, David Spiller, Cyrill Bussy, Kostas Kostarelos, Sandra Vranic

**Affiliations:** Nanomedicine Lab, Faculty of Biology, Medicine and Health, The University of Manchester, AV Hill Building, Manchester M13 9PT, U.K; National Graphene Institute, The University of Manchester, Booth Street East, Manchester M13 9PL, U.K; Division of Neuroscience and Experimental Psychology, School of Biological Sciences, Faculty of Biology, Medicine and Health, Manchester Academic Health Science Centre, University of Manchester, AV Hill Building, Oxford Road, Manchester, M13 9PT, U.K; Lydia Becker Institute of Immunology and Inflammation, University of Manchester, Manchester M13 9PT, U.K; School of Medicine, College of Health and Medicine, University of Tasmania, Hobart, Tasmania, Australia; FBMH Platform Sciences, Enabling Technologies & Infrastructure, FBMH Research & Innovation, Faculty of Biology, Medicine and Health, The University of Manchester, Michael Smith Building, Manchester M13 9PT, U.K

## Abstract

Graphene oxide (GO) holds great potential for biomedical applications, however fundamental understanding of the way it interacts with biological systems is still lacking even though it is essential for successful clinical translation. In this study, we exploit intrinsic fluorescent properties of thin GO sheets to establish the relationship between lateral dimensions of the material, its cellular uptake mechanisms and intracellular fate over time. Label-free GO with distinct lateral dimensions, small (s-GO) and ultra-small (us-GO) were thoroughly characterised both in water and in biologically relevant cell culture medium. Interactions of the material with a range of non-phagocytic mammalian cell lines (BEAS-2B, NIH/3T3, HaCaT, 293T) were studied using a combination of complementary analytical techniques (confocal microscopy, flow cytometry and TEM). The uptake mechanism was initially interrogated using a range of pharmaceutical inhibitors and validated using polystyrene beads of different diameters (0.1 and 1 μm). Subsequently, RNA-Seq was used to follow the changes in the uptake mechanism used to internalize s-GO flakes over time. Regardless of lateral dimensions, both types of GO were found to interact with the plasma membrane and to be internalized by a panel of cell lines studied. However, s-GO was internalized mainly via macropinocytosis while us-GO was mainly internalized via clathrin- and caveolae-mediated endocytosis. Importantly, we report the shift from macropinocytosis to clathrin-dependent endocytosis in the uptake of s-GO at 24 h, mediated by upregulation of mTORC1/2 pathway. Finally, we show that both s-GO and us-GO terminate in lysosomal compartments for up to 48 h. Our results offer an insight into the mechanism of interaction of GO with non-phagocytic cell lines over time that can be exploited for the design of biomedically-applicable 2D transport systems.

## Introduction

Graphene oxide (GO), the oxidized form of graphene, has been one of the most researched 2-dimensional (2D) materials in nanomedicine due to advantageous intrinsic properties^1–3^. For example, GO has large surface area allowing high capacity for loading of cargos *via* both covalent and non-covalent interactions, while different (carboxyl, epoxy and hydroxyl) functional groups offer chemical routes/anchor sites for further functionalization. Most importantly, GO is amphiphilic material that contains hydrophobic carbon basal plane and hydrophilic oxygen-rich functional groups. This favours cellular attachment and proliferation, but also provides good dispersibility in aqueous environment^4–6^. Some of the most promising applications of GO include the development of biosensors, drug delivery platforms, bio-imaging agents and photodynamic/photothermic therapeutic agents^6–11^.

Even though there is a significant number of publications proposing potential applications of GO in nanomedicine, answers to some of the fundamental questions such as how does GO interact with the cells, and which properties of GO drive its uptake remains elusive. This is relevant since the difference in the uptake mechanism can affect the distribution and intracellular fate of the material, which in return affects the biological activity of the material. Understanding the dynamics of cellular interactions with GO is fundamental for the design of efficient drug/gene delivery systems using the materials. For instance, in drug delivery, cellular uptake of the drug can be enhanced by exploiting receptor-mediated uptake pathways^12^. In gene therapy, which commonly targets cell nucleus/cytoplasm, it is important to avoid lysosomal compartments where enzymatic degradation takes place^13^. Cargos internalized *via* phagocytosis will fuse directly with the lysosome for degradation. In contrast, cargos internalized *via* caveolae-mediated endocytosis can be sent from one endocytic compartment to another, eventually reaching the lysosome or sorted to the trans-Golgi network or to basal side of the cell *via* transcytosis^14^. Moreover, questions concerning the toxicological profile of GO cannot be fully answered without knowing if the material can be taken up by the cells: plasma membrane disruption and/or internalization of the material might be the mechanisms leading to adverse effects on the cellular level.

Previous *in-vitro* studies focused on understanding the relationship between intrinsic properties of GO and its cellular internalization. An extensive collection of studies demonstrated that GO could be chemically modified to enhance or reduce its internalization in specific cell lines^9,15–20^. Currently, flow cytometry, confocal laser scanning microscopy (CLSM) and/or fluorescence microscopy have been used as primary tools for assessing cellular interactions with GO. However, this requires the attachment of the fluorescent label to GO through covalent or non-covalent bonding and might alter the intrinsic properties of the material. Furthermore, the inherent differences in starting material, chemical modifications and experimental conditions used have made it difficult to draw general conclusions about interactions of GO with cells. In this regard, a systematic investigation of how unlabeled GO interacts with cells over time, employing a combination of complementary analytical techniques with a thorough characterization of the non-labelled material is urgently needed.

Size of the material is one of the most important parameters which will determine the way that material interacts with cells^21,22^. Previous studies showed that GO with different sizes was taken up *via* different uptake mechanisms^23–25^. To assess the uptake mechanism, most of the studies use pharmacological inhibitors. Important limitation to this approach is inherent toxicity, limiting duration of the study and lack of specificity of the inhibitors used^26^. Therefore in this study, we decided to use label-free non-functionalized GO and investigate the uptake mechanism of GO based on a mix-method approach. This includes the use of pharmacological inhibitors at the early time-point (with a wide range of inhibitors) and expression analysis of genes involved in the regulation of the main uptake pathways by RNA-seq at the later time point to understand changes over time. Furthermore, polystyrene beads of different sizes were used as an additional measure to validate the specificity of the inhibitors. We aim to provide a systematic investigation of the dynamic interactions of GO with the cells, focusing on the dynamic changes of the uptake mechanism, and the intracellular fate of small GO (s-GO, with average lateral size of 477 ± 270.4 nm) and ultrasmall GO (us-GO, with average lateral size of 124.8 ± 69.8 nm) in a panel of cell lines. We hypothesize that GO can be taken up by a range of non-phagocytic cell lines, different flake sizes will be taken up *via* different uptake mechanisms, and main uptake mechanism of GO will change over time.

Four mammalian, non-phagocytic cell lines were selected in this study: human epithelial lung cell line (BEAS-2B), human epithelial skin cell line (HaCaT), human epithelial embryonic kidney cell line (293T) and mouse fibroblast embryonic cell line (NIH/3T3).

Herein, we applied the methodology established in our previous work^27^ allowing realtime observation of the interactions of label-free GO at a single-cell level. We used thoroughly characterized materials differing only in their lateral dimension, a combination of quantitative and qualitative analytical techniques (CLSM, flow cytometry, TEM) to assess the uptake of GO, pharmaceutical inhibitors, RNA-seq and polystyrene microspheres to reveal and validate the uptake mechanism of GO overtime, and lastly live-cell time-lapse imaging to question the intracellular fate of GO.

## Results and Discussion

### Preparation and characterisation of GO

We used GO with two distinct lateral dimensions: s-GO and us-GO, both synthesized following the experimental protocol previously described by Rodrigues *et al*.^4^. The two graphene oxide materials were thoroughly characterised using a range of techniques (**Table 1**).

**Table 1.** Physico-chemical characterization of s-GO and us-GO used in this study.

| Parameter | Technique | sGO | usGO |
| --- | --- | --- | --- |
| Optical properties | Absorbance | $A_{230} = 0.046 \cdot C_{GO}$ (R=0.999) | $A_{230} = 0.048 \cdot C_{GO}$ (R=0.999) |
| | Fluorescence | $E_{595} = 0.73 \cdot C_{GO}$ (R=0.997) | $E_{595} = 0.72 \cdot C_{GO}$ (R=0.996) |
| Degree of defects ($I_D/I_G$) | Raman | $1.2 \pm 0.01$ | $1.2 \pm 0.03$ |
| Zeta potential | ELS | $-62 \pm 1.3$ mV | $-58 \pm 1.3$ mV |
| Oxidation degree | TGA | 43.1 % | 43.2 % |
| Surface chemistry:<br>• Chemical purity<br>• C:O ratio<br>• $sp^2/sp^3$<br>• C-O<br>• C=O<br>• O-C=O<br>• $\pi-\pi^*$ | XPS | 99.6 ± 0.09 %<br>2.4 ± 0.1<br>44.4 ± 3.1<br>45.3 ± 3.3<br>3.2 ± 0.5<br>5.0 ± 0.9<br>1.9 ± 0.6 | 98.9 ± 0.29 %<br>2.2 ± 0.1<br>46.9 ± 3.3<br>42.6 ± 1.8<br>1.3 ± 1.1<br>6.5 ± 0.6<br>2.4 ± 1.1 |
| Lateral dimension (size)* | AFM | 20 – 800 nm<br>(average: 91.2 ± 81.3 nm, 82 % ≤ 125 nm, 8 % ≤ 225 – 150 nm) | 20 – 500 nm<br>(average: 70.9 ± 46.2 nm, 90 % ≤ 125 nm) |
| | TEM | 0.1 – 1.6 μm<br>(average: 477.2 ± 270 nm, 14 % ≤ 225 nm, 76 % ≤ 875 – 250 nm) | 30 – 480 nm<br>(average: 124.8 ± 69.8 nm, 90 % ≤ 225 nm) |
| Thickness | AFM | 94.4 % ≤ 2 nm | 99.19 % ≤ 2 nm |

\* The lateral dimension (size) range was reported as the smallest and highest flake size measured by AFM and TEM. ELS = electrophoretic light scattering, TGA = thermogravimetric analysis, XPS = X-ray photoelectron spectroscopy, AFM = Atomic force microscopy, TEM = Transmission electron microscopy, $A_{230}$ = Absorbance at 230 nm, $E_{595}$ = Emission at 595 nm, $C_{GO}$ = Graphene oxide concentration.

Optical properties of both GO suspensions were evaluated by UV-vis and fluorescence spectroscopy. Both s-GO and us-GO were found to display an intense UV-vis absorbance peak at 230 nm, while an emission band located at 595 nm dominated their fluorescence spectra. The intrinsic fluorescent properties of GO were investigated further using fluorescence spectroscopy: s-GO exhibited higher intensity of auto-fluorescence than us-GO in both water and cell culture medium with serum (**Figure S1**). This is in agreement with our previous finding that auto-fluorescence of GO positively correlates with lateral dimensions of the flakes, i.e. larger GO displays stronger auto-fluorescence comparing to the smaller GO^27^.

Structural characterisation of GO was analysed by Raman spectroscopy, showing the presence of defects in the sp^2^ backbone (I_D_/I_G_ = 1.2) due to incorporation of oxygen functional groups during the synthesis. These oxygen species were responsible for good colloidal stability of the material, as also reflected by the Zeta potential values below −30 mV. Surface chemical characterisation was analysed by X-ray photoelectron spectroscopy (XPS). Elemental analyses showed a very low degree of impurities (≤ 1%) after the oxidation treatment and approximately 30% oxygen content, as indicated by carbon to oxygen (C:O) ratios of 2.2 and 2.4 for s-GO and us-GO, respectively. Additionally, both s-GO and us-GO present similar percentages of oxygen functional groups obtained from the deconvolution of the C1s XPS spectra.

Finally, the difference in lateral size between s-GO and us-GO was a result of different sonication times applied to the starting GO dispersions. Interestingly, the lateral size distribution measured by AFM and TEM showed slight differences, for example, s-GO (and us-GO) had a size distribution between 20 – 800 nm (20 – 500 nm) when AFM was used, but 0.1 – 1.6 μm (30 – 480 nm) when measured by TEM. The average size of s-GO (us-GO) is 91 ± 81 nm (70.9 ± 46.2 nm) by AFM measurement, and 477.2 ± 270 nm (124.8 ± 69.8 nm) by TEM measurement. The differences in lateral size distributions obtained by AFM and TEM were mainly due to a difference in image contrast and resolution between the two techniques. On the one hand, we fixed the AFM scan size to 5 μm in order to have a good view of the small size flake population for both s-GO and us-GO. Larger scan sizes would have led to a poorer view of these flakes, *i.e*. with the detected signal similar to noise. On the other hand, the poor contrast of the flakes in TEM led to a better overview of the larger size population compared to very small flakes. For this reason, when reporting the lateral size range of our GO, we refer to the interval between the minimum size detected by AFM and the maximum size detected by TEM. TEM and AFM micrographs presented in **Figure 1** (**A** - **B** and **D** - **E**) show thin and differently sized s-GO and us-GO flakes. Statistical analysis (**Table 1**) of the flakes showed that the two sized GO have a similar lateral size distribution for the small flakes population (measured by AFM), but the two materials have significant lateral size differences for the large flakes population. As indicated in **Table 1**, the statistical analysis of the flakes measured by TEM showed that 90% of the us-GO flakes had lateral sizes of 225 nm and below. In contrast, only 14% of the s-GO flakes had lateral sizes of 225 nm and below, 76% of the s-GO flakes had lateral sizes of 875 nm and below but greater than 225 nm. In addition, AFM also indicates that GO consists of mainly monolayer and bilayer graphene oxide flakes, i.e. 94% of s-GO and 99% us-GO flakes have a thickness of below 2 nm (**Table 1**). In summary, our results demonstrate that the physicochemical properties and the thickness of both materials are preserved throughout the synthesis process, even though the sizes of the flakes were different.

**Figure 1.**
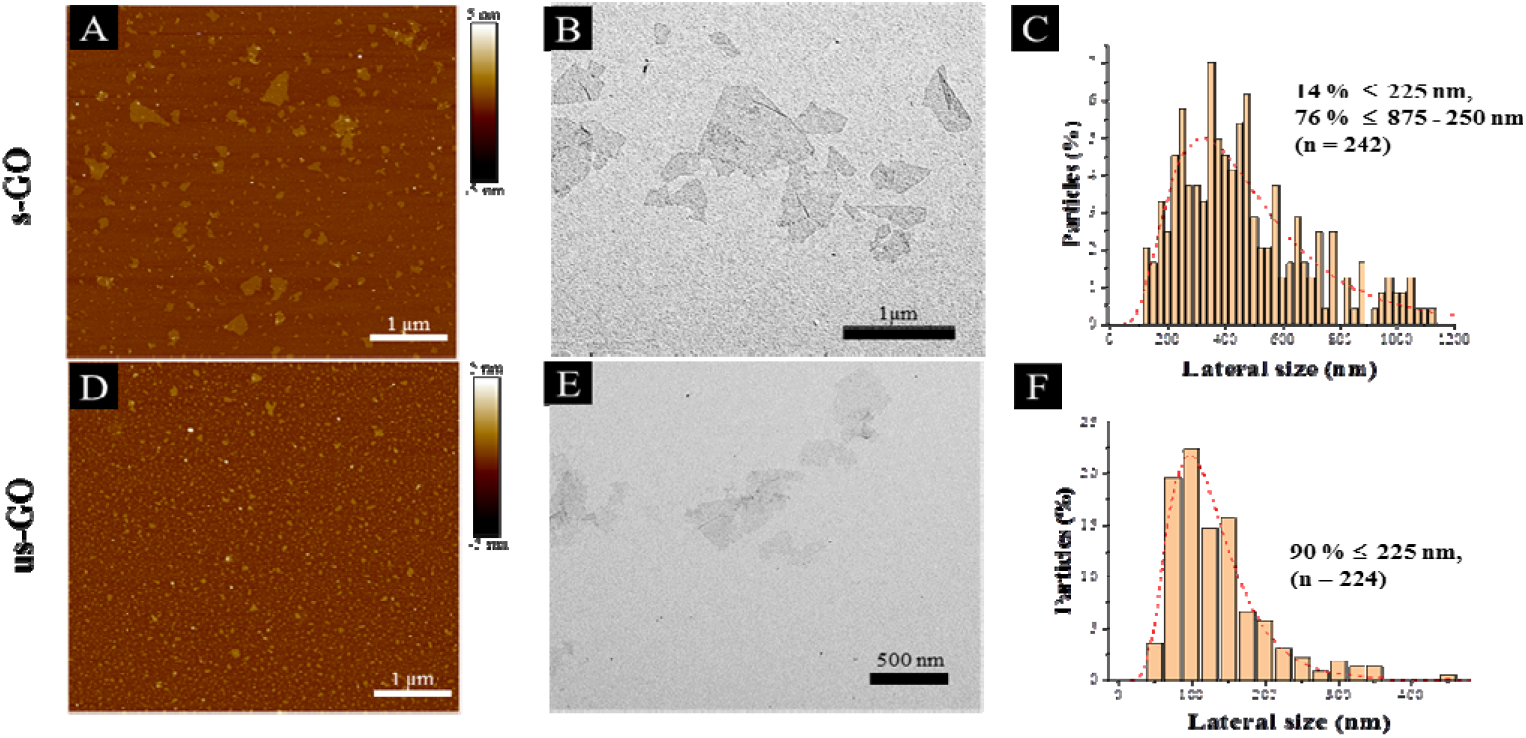
Physico-chemical characterization of s-GO (**A-C**) and us-GO (**D-F**): AFM profile images (**A** and **D**), TEM images (**B** and **E**) and the lateral size distributions determined from TEM images (**C** and **F**).

### Characterisation of GO in cell culture medium

Another aspect to take into account when studying interactions of GO with cells is the new identity material obtains in the biological environment. For instance, when GO, or any other nanomaterial is dispersed in complete cell culture medium (i.e. which contains serum and other supplements), the surface of GO is instantly covered with various serum proteins, electrolytes and biomolecules which alter the surface properties, colloidal stability and the size of the materials, and hence the way it interacts with the cells^21,28–30^. Few researchers have attempted to evaluate the impact of serum on cellular interactions with GO. For example, Duan *et al*. found that GO coated with serum proteins have lower membrane penetration ability than GO without protein coating^31^, and in our previous study, we found that the presence of serum can alleviate the toxic response induced by large GO flakes^27^. Therefore, with consideration to our previous findings and to mimic the biologically relevant environment the material is exposed to, all experiments in this study were performed in a serum containing environment. Even though lung epithelial cell line (BEAS-2B) was chosen, it is still clinically relevant to perform the study in the presence of serum: the epithelial lining fluid, which is a fluid layer covering the alveolar epithelium, contains some of the most abundant proteins usually found in serum and blood plasma (e.g. albumin, immunoglobulin G and transferrin)^32–34^.

Currently, when it comes to connecting the size of GO with the observed uptake profile, most studies only report the primary size of GO used in the study (*i.e*. lateral dimensions of GO flakes measured by TEM or AFM)^17,35,36^. Moreover, very often it is not specified whether the treatment using GO dispersed in cell culture medium contains serum proteins or not, and does not interrogate whether the colloidal stability of the material is affected^37,38^. As already discussed, materials gain a new identity when dispersed in the biological medium, and hence characterisation of the material in the cell culture medium is extremely important. So far, characterisation of the material in a biological dispersion is often done for material with a spherical geometry but much less considered for 2-dimensional (2 D) material^39,40^. Of the limited studies, characterisation of GO in the cell culture medium has been performed by incubating the material in the cell culture medium, followed by material isolation and characterisation by AFM^24,31^.

To date, only a few studies have reported *in situ* characterisation of GO in liquids, mainly by light scattering techniques^28,41–43^. In contrast to TEM and AFM in the dry state, light scattering techniques such as dynamic light scattering (DLS), bring the advantage of a faster and simpler method to evaluate the size of 2D materials dispersed in liquids. Although DLS is based on assumptions for spherical particles, different research groups have shown that the lateral size of 2D materials measured by TEM and AFM were actually scaled reasonably well with measurements by DLS^41,43^. Hence, DLS can be used to offer valuable information on the relative changes in the size of the material in liquid over time. Based on such considerations, we evaluated the change of size and surface charge of s-GO and us-GO upon incubation in RPMI cell culture medium supplemented with foetal bovine serum (RPMI w FBS) for 0 min, 4 and 24 h (**Figure 2**), where the size measurements were done by DLS and Zeta potentials were measured by electrophoretic light scattering (ELS). These time points were selected since interactions of GO with cells were studied after 4 h and 24 h incubation period, and with the extra time point of 0 min, this should give us a better overview of how material changes over time. The changes in the size can reflect the agglomeration/aggregation status of the material, and the Zeta potential can give an indication of the surface charge and colloidal stability of GO in the cell culture medium.

**Figure 2.**
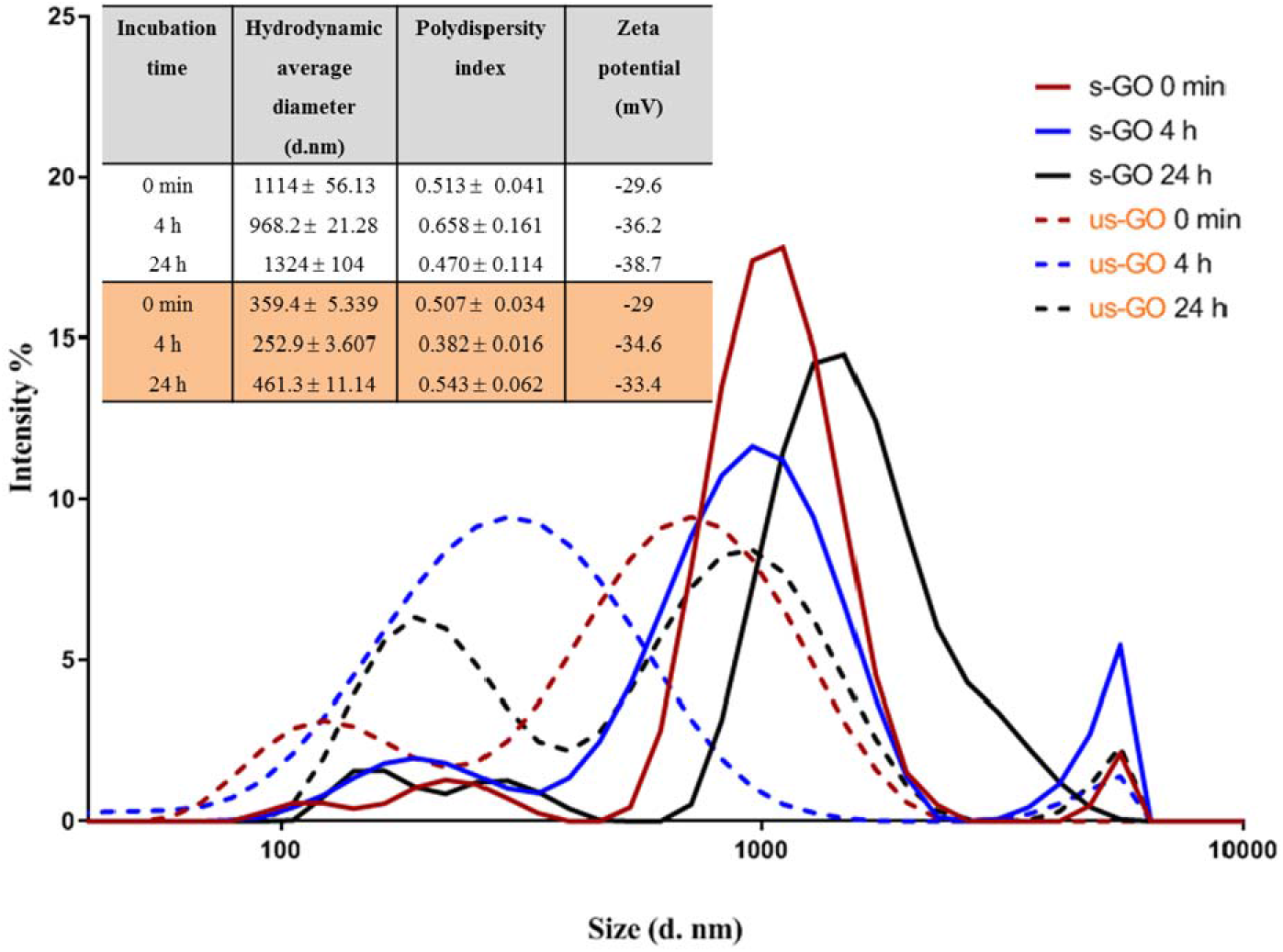
Size distribution of s-GO (solid lines) and us-GO (dotted lines) incubated in RPMI with FBS at different time points. The graph illustrates the relative changes in the size of s-GO and us-GO after incubation in complete cell culture medium for a different period of times. Inserted table shows the measured hydrodynamic diameters, polydispersity index and Zeta potentials of the materials (in white for s-GO and in orange for us-GO).

The result of the Zeta potential and size distribution (expressed by intensity) of s-GO and us-GO at different time points are shown in **Figure 2**. First, the Zeta potential of both materials was found to be similar across all the time points. The Zeta potential changed from around −60 mV in water (**Table 1**) to −29 mV in RPMI w FBS at 0 min, then gradually increased to the distribution of between −33.4 to −38.7 mV for the 4 and 24 h time points. In general, the higher the absolute Zeta potential, the more stable the material dispersions^44^. This result indicates that GO dispersion in water is more stable than in cell culture medium, which is in agreement with the findings from the literature. For example, Ehrenberg *et al*. showed that Zeta potential of all investigated nanoparticles falls into the distribution range of −40 to −20 mV after 2h of incubation in FBS contained solution, irrespective of the initial surface charge (positively or negatively charged nanoparticles)^45^.

In general, the polydispersity index for s-GO is slightly higher than us-GO, which indicates a broader size distribution for s-GO than us-GO. This is in agreements with findings from both TEM and AFM (**Table 1**). What is interesting is that for both materials, the average size at 0 min is greater than the size measured at the 4 h time point, and then increases again at the 24 h time point. The increase in the size of material can take place *via* the irreversible aggregation and/or reversible agglomeration processes; however, DLS does not distinguish between the two processes. A possible explanation for the initial increase in the size of the materials is that the increase of ionic strength in the cell culture medium has resulted in the instant agglomeration of the material, also reflected in the reduced Zeta potential at 0 min. Then the colloidal stability of the materials improved due to coverage of the serum proteins and/or biomolecules, which increases electrostatic/steric repulsion between the materials. Many studies have shown that the presence of serum enhances the colloidal stability of nanomaterial in cell culture medium^46,47^. However, the ultimate stability of the material in suspension depends on the net inter-particles repulsion and attraction forces, and the types of interaction which contribute to the net repulsion/attraction forces can be easily altered with different media composition and materials properties^30^. The increase in the size of GO at the later time point (24 h) are in keeping with the observation that the material sediment in the medium over time. The deposition of s-GO and us-GO on BEAS-2B cells was evaluated at 4 and 24 h using UV-VIS spectrometry (see **Figure S2**). Regardless of the dose used (25 or 50 μg/mL), both materials show similar deposition pattern with around 20 and 50 % of deposition after 4 and 24 h, respectively. Lastly, it is worth mentioning that overlap of the size-distribution between s-GO and us-GO exists for all time points.

### Uptake of GO across a panel of cell lines

Firstly, we wanted to establish whether GO is taken up by mammalian cells used in this study. BEAS-2B cells were treated with both types of GO and incubated for 24 h. CLSM and TEM were used as the primary tool for assessing the uptake and localization of GO in the cells. Cytotoxicity of s-GO and us-GO (0 – 75 μg/mL, 24h treatment) in BEAS-2B cells was assessed *via* optical imaging and Propidium Iodide/Annexin V staining by flow cytometry (**Figure S3**). The result confirmed that neither s- or us-GO induced cell death using a dose range studied. Furthermore, the size-dependent toxicity effect of GO has been thoroughly investigated in our previous work^27^.

Looking at the confocal images of the apical and middle section of the cells in **Figure 3**, we determined the pattern of interactions of the GO with the cells. The internalized GO was easily distinguished from the GO attached on the surface of the cells. The material attached on the surface of the cells appeared as a cloud of signal coming from GO sitting on top of the plasma membrane (**Figure 3A**). On the other hand, the internalized GO (the red-spotted signal) was found distributed in the cytosol and predominately around the nucleus, as indicated by the circular distribution of the material towards the centre of the cells (**Figure 3B**). This observation suggests that the material was intracellularly trafficked towards the lysosomes; upon internalization, the newly formed cargo-enclosed vesicles fuse to form the early endosome, which then migrates from the periphery of the cell to a near-nucleus location where the late endosome will fuse with the lysosome for degradation^48^. The orthogonal projection of the middle section of the cells (**Figure 3C**) confirmed that the circularly distributed material was found inside the cells. Interactions of GO with BEAS-2B cells was also confirmed by TEM (see **Figure 3D**). The TEM images confirmed the results obtained by confocal imaging; both s-and us-GO were found interacting with the plasma membrane and enclosed in vesicles within the cell.

**Figure 3.**
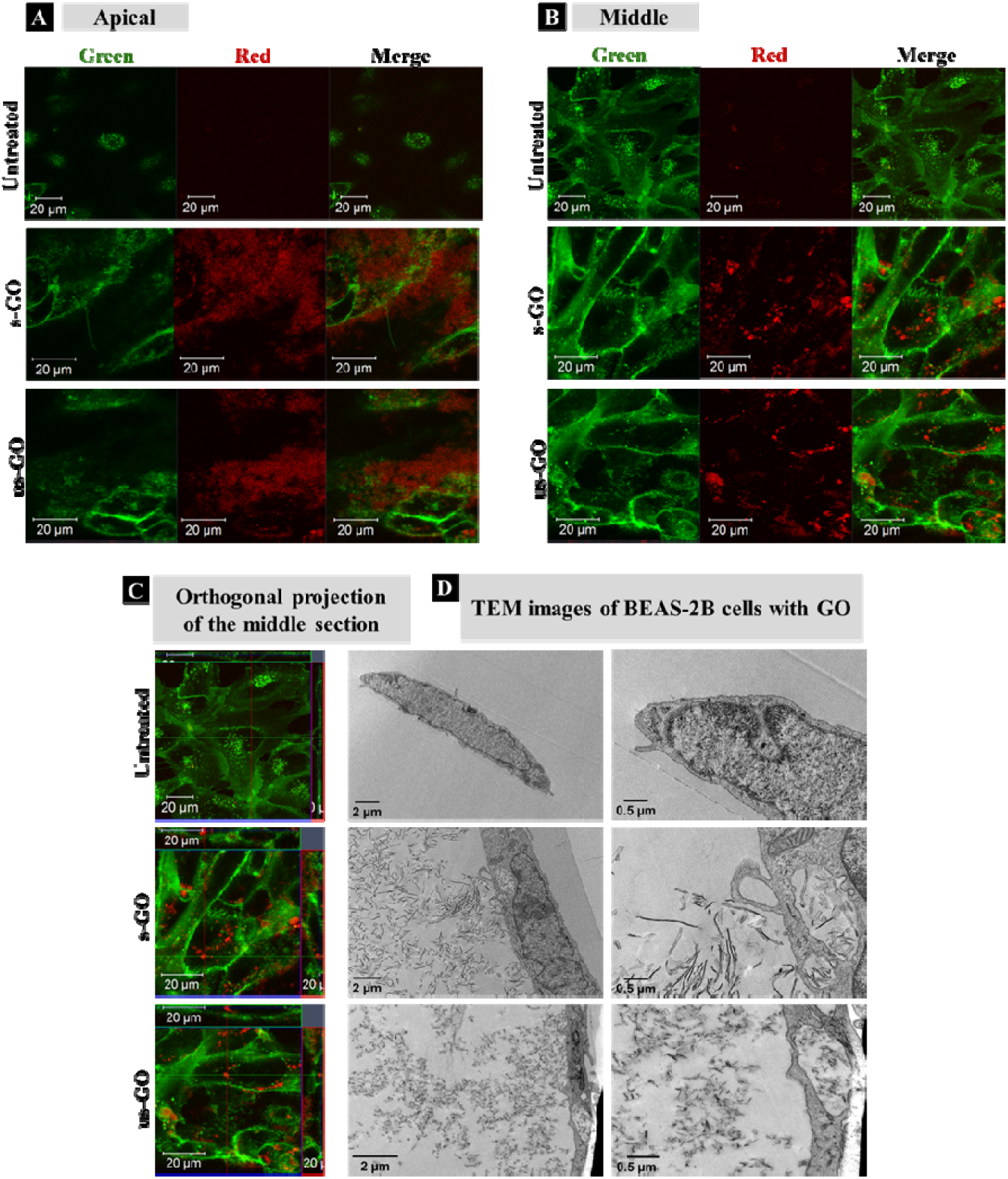
Confocal (**A – C**) and TEM (**D**) images or BEAS-2B treated with 50 μg/mL of GO after 24 h of treatment. (**A**) Apical, (**B**) middle section, (**C**) orthogonal projection of the confocal images. White arrows in (**A**) and blue arrows in (**D**) indicate regions of GO interacting with the plasma membrane of the cells, which differs from internalised GO as shown in (**B**), (**C**), and (**D**) indicated by white rectangles, yellow arrows, and red rectangles, respectively. Green = plasma membrane, Red = GO.

Further analysis using CLSM showed that both s-GO and us-GO were internalized by other non-phagocytic cell lines as well (NIH/3T3, HaCaT and 294T). Interestingly, different cell lines were capable of taking up GO to a different extent (**Figure 4**). NIH/3T3 - mouse embryonic fibroblasts displayed the highest uptake of GO when compared to HaCaT and 293T, which are human skin and kidney cells, respectively. Biological response to particular materials can vary depending on the origin of the cell line used, and indeed it has been previously reported that the interaction of GO with cells was cell-type specific^25,49^. For example, a study reported by Linares *et al*. showed that human bone cancer cells had the highest total interaction with GO, followed by human liver cell and immune cell lines^25^. However, a more profound understanding of what drives this cell-type specificity towards the GO is still missing. Our future research will investigate further into the factors that drive the cell-type specificity toward the GO.

**Figure 4.**
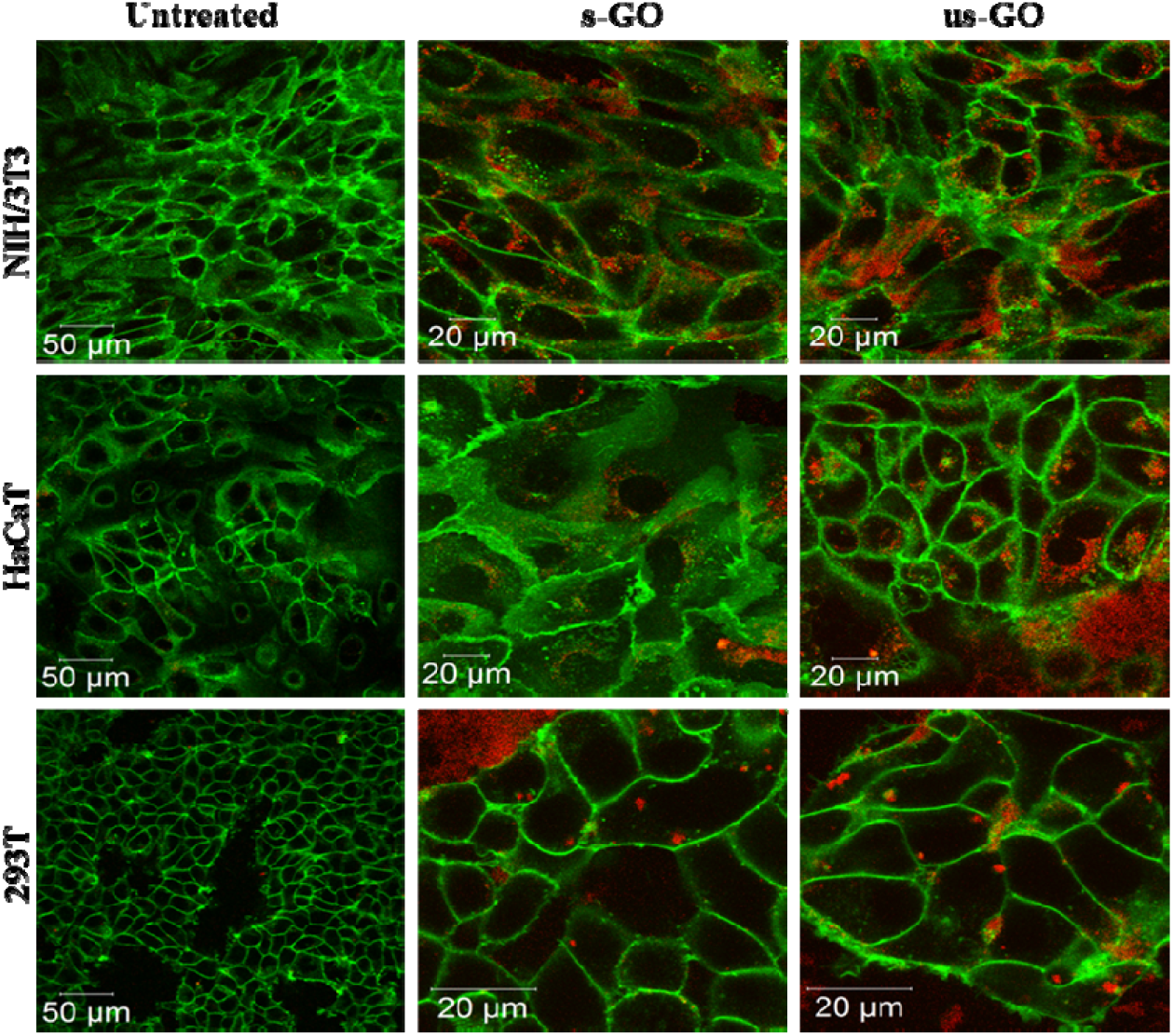
Uptake profile of 60 μg/mL of s-GO and us-GO in NIH/3T3, HaCaT and 293T cells (middle section of the cells is shown) after 24 h of treatment. Green = plasma membrane, Red = GO.

Our results agree with findings from the literature as several studies report internalization of GO by the cells^24,25,27^. However, controversial experimental results are reported as well. For example, Chang *et al*. used GO with lateral size ranging from 160 to 780 nm and observed barely any internalization of GO in human lung carcinoma cells^37^ using TEM, while Jin *et al*., also relied on TEM, reporting an utterly opposite result where GO with lateral dimensions around 300 nm was taken up by the same cell line^35^. Contradictions of the reported findings can occur due to several reasons: such as the batch to batch variation of the GO, the origin of the cell lines and the different treatment conditions of GO, for example, whether treatment was performed in the presence of serum or not. This illustrates the need to include a full characterisation of the materials used, but many of the existing literature on the interactions of GO with cells lack clarity regarding the detailed GO physicochemical properties and experimental conditions used.

Next, we established the time and dose kinetics profile for the uptake of the two sized GO by BEAS-2B cells. Split channel images of BEAS-2B cells incubated for 24 h using different concentrations of s-GO and us-GO (25, 50 and 75 μg/mL) are shown in **Figure 5** (**top panels**). We observe that the two sized GO were successfully taken up by BEAS-2B cells regardless of the size and concentration used, as indicated by the outlined circular distribution patterns of the material. If we compare the uptake of GO prepared in different concentrations, it is clear that its uptake increases with the concentration applied, but no apparent difference observed between the two types of GO. The quantitative analysis of the images (**Figure S4**) confirmed that the uptake of GO in all the treated cells are statistically more significant than untreated cells. The result also confirmed that the cells were taken up GO in a dose-dependent manner. And finally, the result showed no statistical differences in the uptake of s-GO and us-GO by BEAS-2B cells except in the highest concentration of 75 μg/mL.

**Figure 5.**
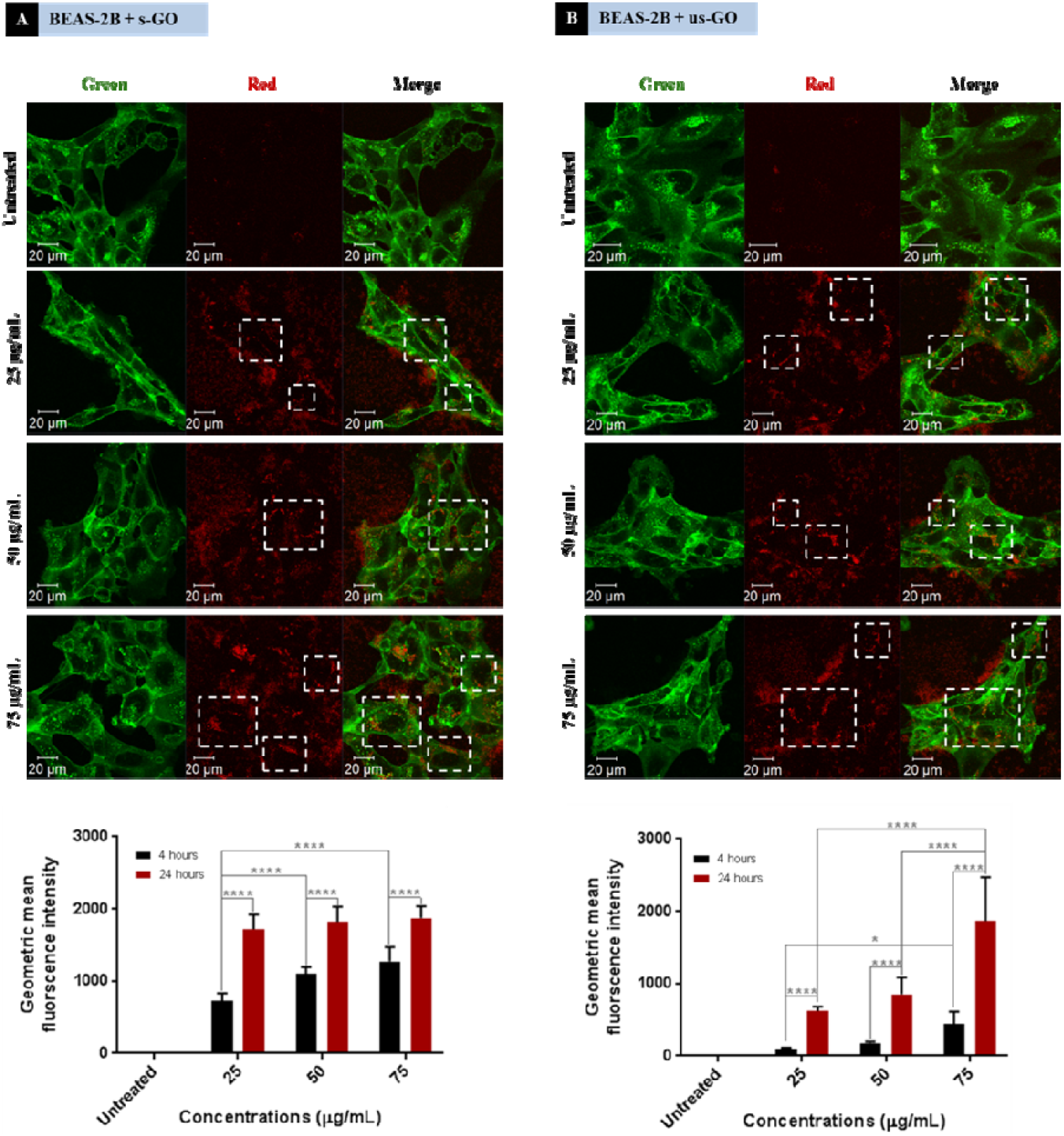
Interactions of (**A**) s-GO and (**B**) us-GO with BEAS-2B cells: (top) dose-dependence of the uptake of GO studied after 24 h of treatment by CLSM, (bottom) dose- and time-dependence interaction of GO with cells studied by flow cytometry. White rectangles indicate the GO. Green = plasma membrane, Red = GO. Flow cytometry data were statistically analysed using analysis of variance (two-way ANOVA) with *post hoc* Sidak’s multiple comparisons test. Difference between treatments at specified concentration and time points with *p* < 0.05 were considered statistically significant \**p* < 0.05 and \*\*\*\**p* < 0.0001. n = 3 independent experiments (each condition in each experiment was run in triplicate).

We further investigated interactions of GO with cells using flow cytometry (**Figure 5** bottom panels). Density plots from flow cytometry are shown in **Figure S5**. Both s-GO and us-GO were found to interact with BEAS-2B cells in a dose- and time-dependent manner, corroborating the results obtained using confocal microscopy. The geometric mean fluorescence intensity (Geo MFI) of the cells was significantly lower at 4 h of incubation compared to 24 h, suggesting time dependence of the interaction. In general, the Geo MFI of the cells treated with s-GO was more intense compared to the ones treated with us-GO. However, the higher intensity measured for s-GO does not necessarily mean that the uptake of s-GO is higher than us-GO. This is because the spectrofluorometric analysis of the materials showed that the s-GO has higher intrinsic fluorescence intensity than us-GO (**Figure S1**). The lack of standardised measure for the quantification of GO flakes in the initial treatment solution has made it challenging to normalise the measured intensity against the quantity of the GO, and so was to assess the size-dependent uptake efficiency of GO. However, when combined with the results obtained from confocal imaging (**Figure 5** top panels), we were able to conclude that the difference in the measured Geo MFI signal between s-GO and us-GO by flow cytometry is most likely due to difference in the intensity of the auto-fluorescent signal between the two materials.

For s-GO, the dose-dependent behaviour was more significant for the 4 h incubation period but showed no significant difference for the 24 h incubation period. In opposite, the dose-dependent response for us-GO was more significant for the 24 h incubation period than the 4 h incubation period. These results indicate a possible difference in the uptake kinetics for the two types of GO. This opens up a new question of whether s-GO and us-GO are taken up by the cells *via* different uptake mechanisms.

### Revealing the cellular uptake mechanism of GO sheets

Eight pharmacological inhibitors for the three main endocytic pathways, including macropinocytosis, clathrin-mediated endocytosis (CME) and caveolae-mediated endocytosis (CavME), were selected to assess the uptake mechanism of s-GO and us-GO in BEAS-2B cell line. Based on flow cytometry, the uptake mechanism of GO was assessed by comparing the fluorescent signal of GO detected in the inhibitors treated cells and cells treated with no inhibitor. It is important to clarify here although flow cytometry cannot distinguish extracellular bound and internalized material, the combination of inhibitor and flow cytometry is the standard technique used in the field to assess the uptake mechanism of cargos. Hence in this section of the paper, we will refer the fluorescent signal detected for the interaction of GO by flow cytometry to the uptake of GO.

Macropinocytosis is involved in the internalization of large particles, ranging in size between 0.2 and 10 μm, which occurs either in an inducible or constitutive manner^48,50,51^. The latter is present in all animal cells, whereas the former takes place mostly in macrophages and dendritic cells. CME takes up specific cargos with the size of around 200 nm^48,50,52–54^. Upon binding of the cargo to a specific transmembrane receptor, a sequence of events stimulated and eventually the coating protein (clathrin) on the cytosolic side of the plasma membrane self-assembles into a cage that forms a coating layer to the invaginated segment of the plasma membrane containing the cargo. And CavME, which is similar to CME, differed mainly in the main coating protein and occurred at plasma membrane region of high lipid content. CavME commonly recognized to take up cargos of a smaller size (~50 to 100 nm)^48,50,55,56^.

**Table 2** summarizes the inhibitors we used, the corresponding affected uptake mechanisms and working concentrations. Series of optimization studies were performed in order to determine the working concentration for each inhibitor in an incubation period of 4 h 30 min^57,58^. The incubation time was selected compromising both the fluorescent intensity of GO that can be detected by flow cytometry and the toxicity of the inhibitor: the incubation period should be long enough to enable the detection of the GO signal, but not too long to induce cell death/stress. As we demonstrated that flow cytometry could be used to detect the signal from GO after 4 h of incubation, this time point was selected. A range of concentrations was tested for each inhibitor (**Figure S6**), and the working concentration was selected based on the principle of selecting the concentration with the maximum inhibitory effect but minimum cytotoxicity induced by the inherent toxicity of the inhibitors^57,59^. The cytotoxicity of the inhibitors was assessed by optical imaging of the cell morphology, propidium iodide (PI)/annexin V (AV) staining using flow cytometry, and the assessment of actin filament disruption using CLSM (**Figure S6 and S7**). Disruption to the actin filaments was assessed to reassure the specificity of the inhibitors as they are involved in several uptake pathways.

**Table 2.**
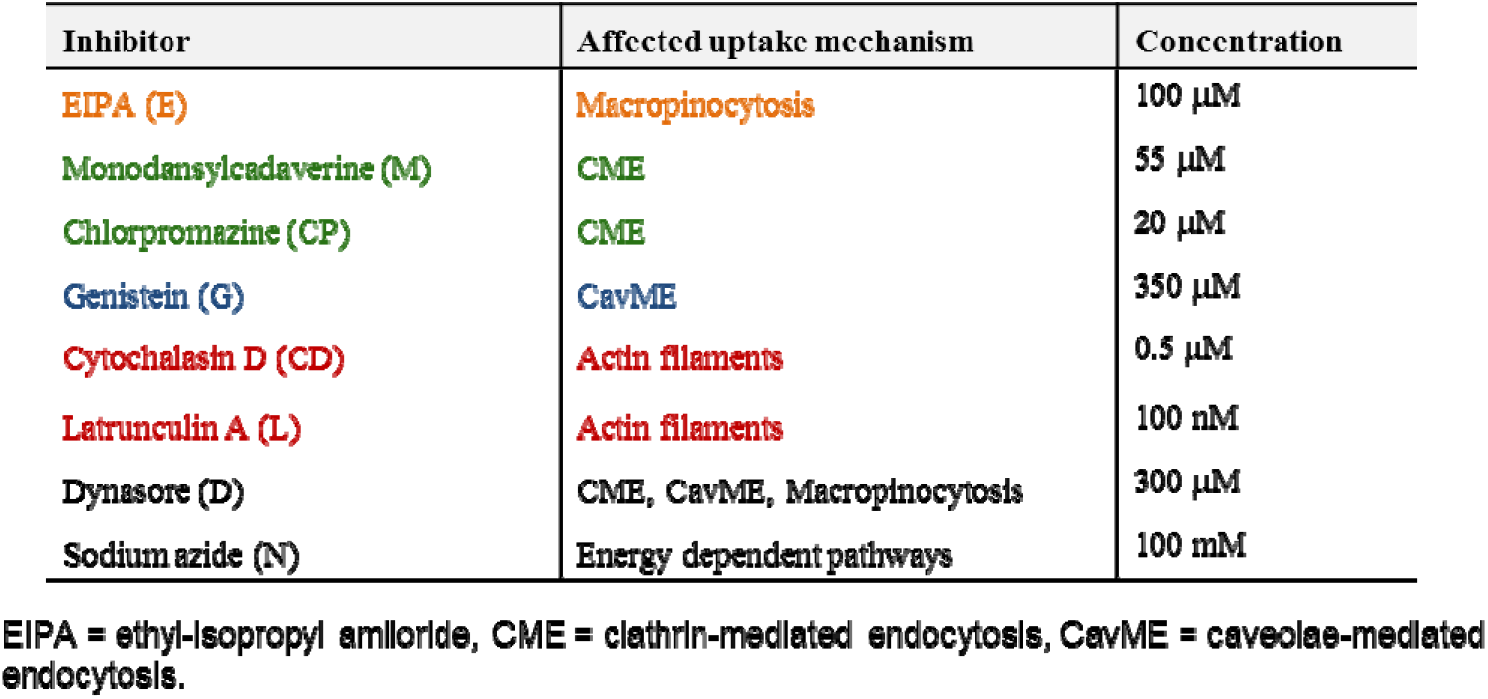
List of the inhibitors used in the study.

In **Figure 6**, we summarize the percentage of inhibition for the uptake of GO in BEAS-2B cell with the treatment of inhibitors. The results were normalized against the corresponding control with no inhibitors. It is important to note that both sodium azide and dynasore were used as non-selective inhibitors for all three uptake pathways described above. Sodium azide affects energy-dependent pathways by inhibiting the process of mitochondria respiration that is responsible for the production of cellular energy. And dynasore inhibits the activity of dynamin which is required for the cleavage of endocytic vesicles, and also known to interfere with actin actin filaments^60^.

**Figure 6.**
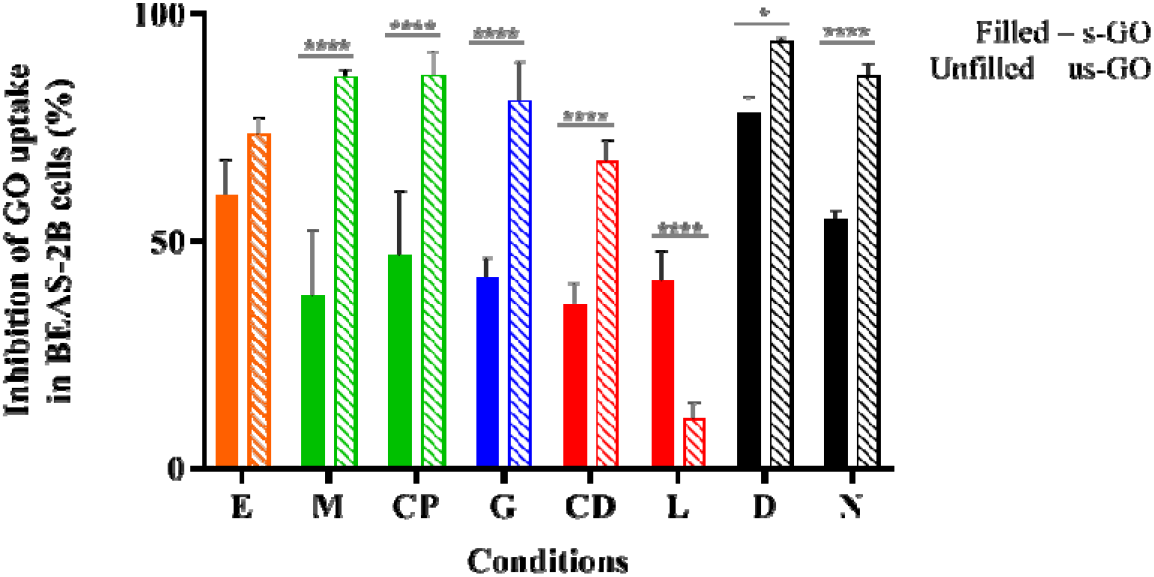
The percentage of uptake inhibition of s-GO (filled) and us-GO (unfilled) in BEAS-2B cells after treatment with inhibitors (E = EIPA, M = Monodansylcadaverine, CP = Chlorpromazine, G = Genistein, CD = Cytochalasin D. L = Latrunculin A, D = Dynasore, N = Sodium azide), assessed by flow cytometry. The data were statistically analysed using analysis of variance (two-way ANOVA) with *post hoc* Sidak’s multiple comparisons test. Difference between treatments with *p* < 0.05 were considered statistically significant: \**p* < 0.05 and \*\*\*\**p* < 0.0001. n = 3 independent experiments (each condition in each experiment was run in triplicates).

As shown in **Figure 6**, dynasore has the highest inhibitory effect on the uptake of GO, but surprisingly, sodium azide was found to be not as effective as dynasore, especially for s-GO. For example, 77.95 and 93.97 % of s-GO and us-GO interaction was inhibited using dynasore, but only 54.80 and 86.22 % of s-GO and us-GO was inhibited with sodium azide, respectively. The significant difference in the uptake inhibition found between the s-GO and us-GO after treatment with sodium azide is rather interesting. A possible explanation is that GO can enter the cells passively, and such process is size-dependent. So far, variety of nanoparticles of different physico-chemical properties (including size, shape, surface charge and functionalization) have previously reported to enters cells *via* non-endocytic pathways, but still very little is known about the factors which drive the process^61–64^. Another possible explanation for the difference between s-GO and us-GO uptake upon treatment with sodium azide is that the main uptake mechanism explored by s-GO can also utilize the metabolic energy supplied by another source, such as glycolysis. It is known that the metabolic energy needed for endocytosis can be derived from glycolysis and respiration, and the dependence on the metabolic energy source varies with cargos and cell types^65^. For example, Steinman et al. (1974) observed the inhibition of horseradish peroxidase uptake in mouse fibroblast is greatly enhanced with combination use of glycolytic and respiratory inhibitors (such as 2-deoxyglucose and sodium azide, respectively); the individual use of glycolytic or respiratory inhibitor resulted in only partial inhibitory effect^66^. In contrast, the uptake of transferrin in protozoan is significantly inhibited with solely the use of 2-deoxyglucose or sodium azide^67^. The literature on GO uptake mechanism in mammalian cells has indicated that the combination use of glycolytic and respiratory inhibitors to exhibit a greater inhibitory effect on the uptake of GO than using single metabolic energy inhibitor. For example, Mu *et al*. found more than 80% of both large and small GO (with the lateral dimension of 1 μm and 500 nm, respectively) uptake is inhibited with the combination of sodium azide and 2-deoxyglucose^24^. In comparison, using gold nanoparticles loaded GO with the lateral dimension of 100 - 200 nm, Huang *et al*. detected the GO signal reduced from 38.5 to 17.4 % with the pre-treatment of sodium azide, which equivalent to only 54.8 % of inhibition^23^.

From **Figure 6**, it is apparent that all the inhibitors had an effect on both s-GO and us-GO uptake, but to a different extent. This suggests that all three main pathways are involved in the uptake of both materials, which is not a surprise, considering the broad size distribution of the two materials. Overall, the inhibitors were found more effective at inhibiting the uptake of us-GO then s-GO. This supports findings from the literature that smaller GO flakes are in general taken up more efficiently than GO of a bigger size^68^. We found that the primary uptake pathway for s-GO is macropinocytosis as ethyl-isopropyl amiloride (EIPA) which affect macropinocytosis, was most effective at inhibiting the uptake of s-GO (60.01 % of uptake inhibition). However, knowing that macropinocytosis is an actin-dependent process, it was surprising that Cytochalasin D (CytD) and Latrunculin A (LA), which perturb polymerization of the actin filaments (F-actin), were not as effective as expected (36.32 and 41.11 % of inhibition respectively)^69,70^. A possible explanation might be the fact that EIPA works by inhibiting the exchange of sodium/proton cation at the surface of the plasma membrane and is therefore more efficient than CytD and LA which affects the actin filaments underneath the plasma membrane^71,72^.

The inhibitors responsible for CME (monodansylcadaverine and chlorpromazine), were the most efficient at inhibiting the uptake of us-GO (with more than 86 % of uptake inhibition for both inhibitors). And genistein, which inhibits the CavME, was effective at inhibiting 80 % of the uptake for us-GO but just over 42 % for s-GO. It is also worth noting that CytD was capable of inhibiting 67.47 % of the us-GO, but LA only reduced the uptake of us-GO by 10.90 %. This result is consistent to Fujimoto *et al*., who found CytD inhibited the uptake of transferrin by 60 – 70 % while LA reduced the uptake by small percentage^70^. Discrepancies in the effect of CytD and LA were found highly dependent on the cell types and the cell adhesion properties^70^.

Overall, results on the involvement of different pathways for the two materials seem to follow the rough guideline of endocytosis determined by the size of the material. As illustrated in **Figure 2**, the size average of s-GO and us-GO incubated in cell culture medium for 4 h was 968.2 ± 21.28 nm and 252.9 ± 3.607 nm, respectively. These results agree with literature findings that macropinocytosis is involved in the uptake of large material (0.2 – 10 μm) whereas CME is commonly known to take up materials of around 200 nm^48,50–54^. Overlap of the size distributions between s-GO and us-GO explained the reason why multiple uptake pathways were involved for both materials.

### Validation of the uptake mechanism

Taking into consideration that there is a growing concern around specificity and efficacy of inhibitors towards one or another endocytic pathway^25,69,73^, we decided to use fluorescently labelled carboxylate-modified polystyrene beads with different sizes (0.1 and 1 μm) to validate the endocytic pathway that we found to be predominantly used by s-GO or us-GO. The beads were selected because both the beads and GO have a similar surface charge (**Table S1**) and chemical composition (GO also contained carboxylate functional groups). Since the size of the cargo can indicate the endocytic pathway involved, 1 μm beads are expected to be internalized mainly *via* macropinocytosis while 0.1 μm beads can be internalized *via* both CME and CavME^58,74,75^. Therefore, we hypothesized that if our findings on endocytic pathways used by GO were true (**Figure 6**), then, the pre-treatment of s-GO or us-GO should reduce the uptake of 0.1 and/or 1 μm beads by occupying or saturating the corresponding pathway. With the broad size distribution of s-GO, we expected the uptake of s-GO to influence the uptake of both 1 and 0.1 μm beads, whereas the uptake of us-GO to affect only the uptake of 0.1 μm beads.

As shown in **Figure 7A** (and corresponded density plots in **Figure S8**), s-GO was very efficient at reducing the uptake of the beads irrespective of their size, while us-GO was efficient at reducing the uptake of the 0.1 μm beads only. For example, 76.1 % (3.0 %) and 77.9 % (49.4 %) of 1 and 0.1 μm beads uptake was inhibited by s-GO (us-GO), respectively. Looking at the confocal images from **Figure 7B-C**, the results agreed with findings obtained using flow cytometry: the uptake of 1 μm beads was significantly reduced with the pretreatment of s-GO but not us-GO. Our previous confocal images indicated the association of GO with the plasma membrane, so we wanted to clarify that the inhibition of the uptake of the beads was not due to the attachment of GO to the plasma membrane and therefore shielding the cells from being in contact with the beads.

**Figure 7.**
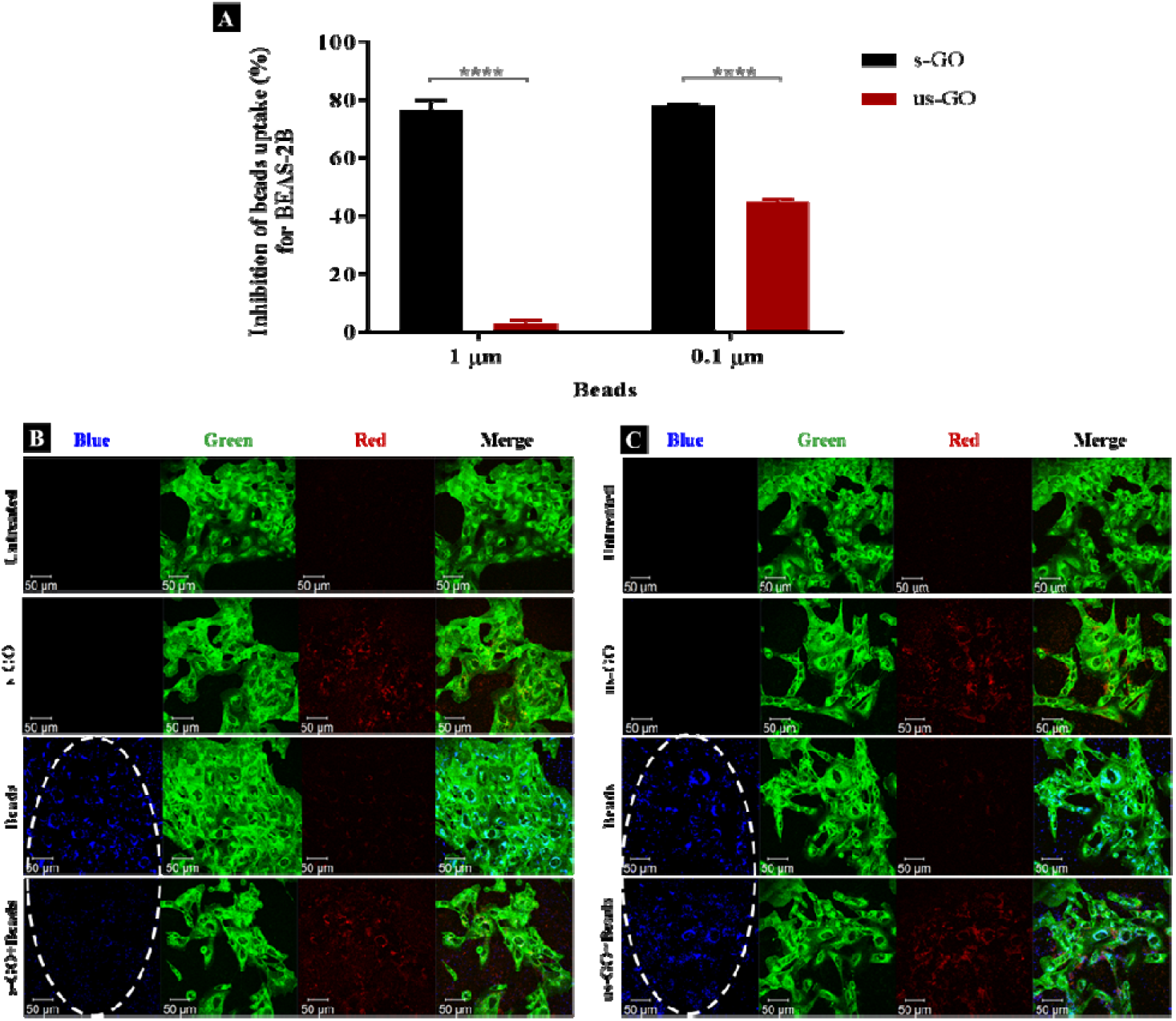
(**A**) Percentage of inhibition on the uptake of beads (1 and 0.1 μm) in the presence of s-GO (black) or us-GO (red), assessed by flow cytometry. The data were statistically analysed using analysis of variance (two-way ANOVA) with *post hoc* Sidak’s multiple comparisons test Difference between treatments with *p* < 0.05 were considered statistically significant \*\*\*\**p* < 0.0001. n = 2 or 3 independent experiments (each condition run in quintuplicates or duplicate, respectively). (**B-C**) The uptake of 1 μm beads with and without s-GO (**B**) or us-GO (**C**), assessed by confocal microscopy. The uptake of the beads was inhibited in the presence of s-GO, but not us-GO, indicated by the white ovals. Green = plasma membrane, Red = GO, Blue = 1 μm beads.

Looking at the apical section of the cells (**Figure S9**), cells treated with us-GO and 1 μm beads showed an obvious co-localisation of the two materials on top of the plasma membrane. However, this was not the case for cells treated with s-GO and 1 μm beads (**Figure S9**); instead, clouds of s-GO signal without obvious clustering with the beads were found. Considering that the treatment of us-GO did not prevent the uptake of the beads, we conclude that the reduction in the uptake of beads was not due to the shielding effect of the GO but rather to the interference of GO with the corresponding uptake pathway.

In summary, our results show that all three major uptake pathways were involved in internalization of s-GO and us-GO by BEAS-2B cell line, but s-GO preferentially used macropinocytosis, while us-GO used mainly CavME and CME. This was possible due to the broad size distribution of GO flakes: the smallest flakes were internalized mainly *via* CavME and CME, while the bigger ones entered the cells mainly *via* macropinocytosis. So far, surprisingly, only a few studies found in the literature focused on understanding GO uptake mechanism by the cells^23–25^. Our results are in agreement with the literature showing that the uptake of GO occurs *via* energy-dependent pathways^23–25^. The discovery of both s-GO and us-GO internalized *via* macropinocytosis was consistent with Linares *et al*. who found macropinocytosis to be an uptake mechanism for FITC-PEG-GO (with the lateral dimension of 100 nm) in the human liver cancer cell (HepG2) and human bone cancer cell (Saos-2)^25^. Both phagocytic cells (C2C12, Raw-264.7) and non-phagocytic cells (HepG2, HeLa, Ca Ski) were found to take up GO through CME; despite the GO used have different physicochemical properties (with the lateral dimension ranged from 100 nm to 1 μm).

### Temporal change of cellular interactions with s-GO

To track the dynamics of GO uptake over time, RNA-Seq was used to analyse the expression of the genes involved in the main uptake pathways 24 h after the exposure to s-GO. The main limitation of pharmacological inhibitors is the duration of the treatment that should not exceed 6 h, due to the abovementioned toxicity and lack of specificity issues. RNA-Seq offers an alternative to interrogate later time points and reveal changes in the expression of genes involved in different uptake pathways. We treated BEAS-2B cells with 50 μg/mL s-GO, and then extracted the RNA for analysis by RNA-Seq (**Figure 8 A-D**). As reported in **Figure 8 A**, the principal component analysis confirmed significant differences between the transcriptomes of the untreated and cells treated with s-GO for 24 h. To probe this difference, enrichment analyses were performed on the significantly upregulated and downregulated genes using an unbiased approach and probing the *apriori* hypothesised pathways of macropinocytosis, CME and CavME (**Figure 8B-D**). These analyses revealed a shift in uptake mechanisms following s-GO exposure with the downregulation of macropinocytosis and upregulation of CME and CavME (**Figure 8B-D**). Furthermore, our enrichment analyses found that mTORC1/2 pathways were highly upregulated (**Figure 8D**). Work by Srivastava et al. 2019 found that mTORC1/2 inhibition promoted macropinocytosis^76^. Combining this research with the result reported here, we propose that 24h exposure to s-GO induces mTORC1/2 activation, which both inhibits macropinocytosis and activates CME and CavME, shifting the pathways of uptake (**Figure 8E**)^77^.

**Figure 8.**
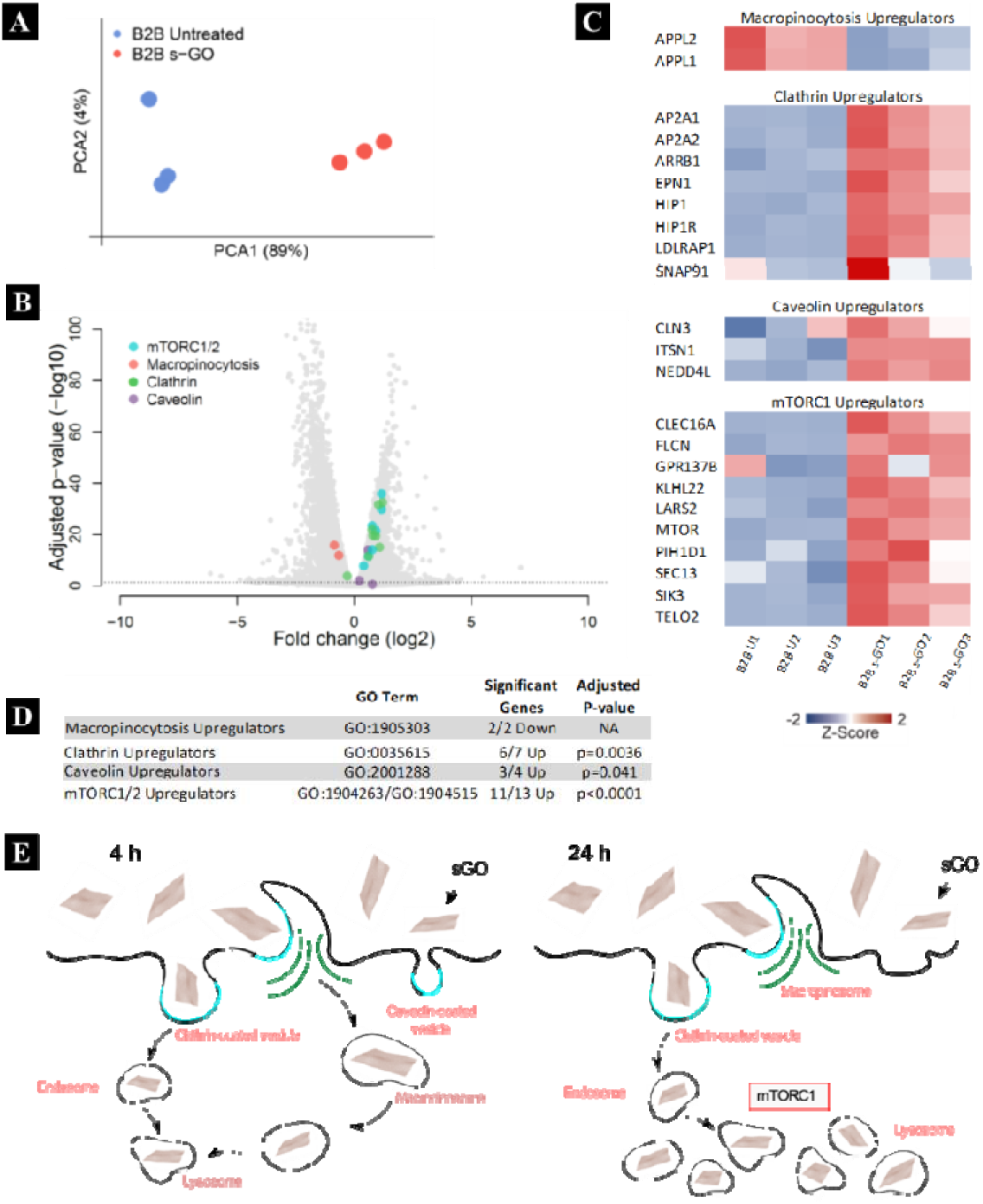
Transcriptomes data showing differential gene expression between untreated and BEAS-2B calls treated with 50 μg/mL s-GO for 24 h. (**A**) Princioal component analysis plot confirme substantial difference between the transcriptomes obtained under the two condition tested. (**B**) Volcano plot indicating that a large number of genes are up- and downregulated 24 h after treatment. (**C**) Heatmap showing differences between the relative expression of genes related to macropinocytosis, clathrin-, caveolae-mediated endocytosis as well as mTORC1 regulation. (**D**) Enrichment analyses showing significant shifts in upregulators of three main endocytic pathways as well as of mTORC1/2. (**E**) Proposed shifts in the uptake mechanism used by BEAS-2B cells at 4 h (left) and 24 h (right), with s-GO being transported to the lysosomes. Due to saturation, upregulation of the mTORC1 complex prevents further macropinocytosis, despite continuing ruffle formation.

### Intracellular fate of GO

We interrogated intracellular fate of GO with the use of CellLight™ Lysosomes-GFP. **Figure 9** displays snapshots of the images taken at 24 h and 48 h time points during live-cell time-lapse experiment (**Video S1-S5**). The images show that both s-GO and us-GO end up in the lysosomes, as indicated by localisation of the GO signal (red) enclosed by the signal of the lysosomal marker (green).

**Figure 9.**
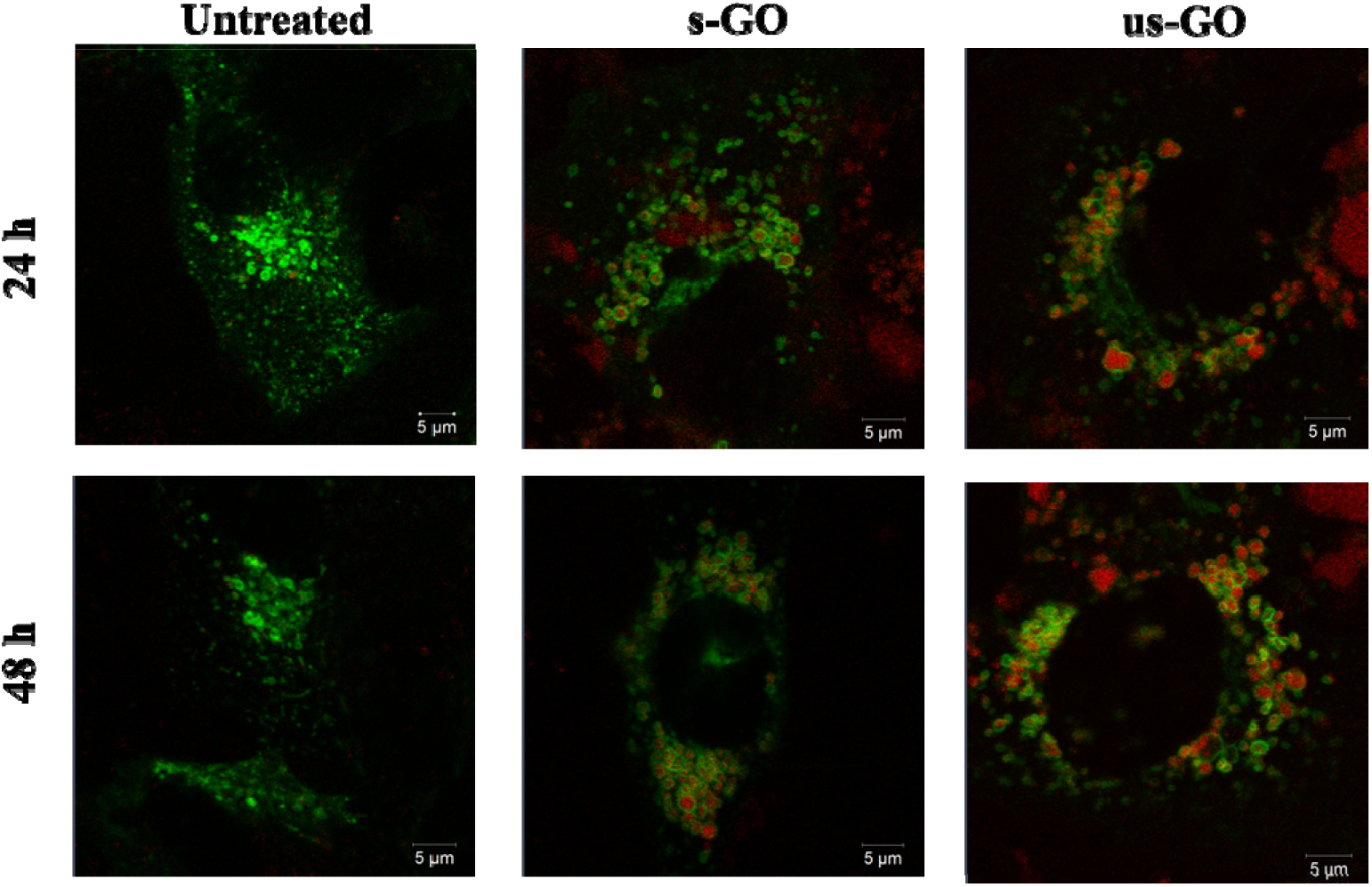
Snapshots of time-lapse videos of the BEAS-2B cells treated with s-GO or us-GO for 24 and 48 h. Lysosomal compartments were labelled using CellLight™ Lysosomes-GFP, BacMam 2.0 prior to the treatment with GO. Green = lysosomes, Red = GO.

This result is in agreement with the literature suggesting that internalized GO ends up in the lysosomes^9,24^. Some studies show that surface charge of GO can be modified using different chemical moieties to influence the intracellular fate of GO^16,18,78^. For example, utilising mouse macrophages, Wang *et al*. found positively charged polyethyleneimine modified GO (PEI-GO) in both endosomes and cytoplasm, while negatively charged polyethylene glycol-amine modified GO found only in the endosomes^18^. Tripathi *et al*. demonstrated that the positively charged linear PEI-GO end up in the nucleus of cancer and non-cancer cells^16^. And Jin *et al*., who also used cancer cells, reported long term retention of the positively charged PEI-GO in the mitochondria, which could be due to the leakage of PEI-GO into the cytoplasm^78^. Therefore, even though the majority of GO was found in the lysosomes in this study, knowing the limitations of analysing non-labelled materials, we do not rule out the possibility that some individualized flakes might end up in the cytoplasm of the cells.

From **Video S1-S5**, we could also see a consistent exchange of both materials between the lysosomal vesicles, and with no obvious disruption to the lysosome membrane up to 48 h of treatment. These findings confirm results from our previous study, where we observed no toxicity or lysosomal rupture after treatment with s-GO using cell culture medium supplemented with 10% FBS^27^. Based on the results of this study, we can extend this statement to us-GO as well.

## Conclusion

This study aimed to provide a better understanding of the fundamental interactions of label-free GO with non-phagocytic mammalian cells (BEAS-2B, NIH/3T3, HaCaT and 293T) by taking advantage of the intrinsic fluorescent properties of GO. GO with two distinct lateral dimensions were used in the study. Using confocal microscopy, flow cytometry, TEM and RNA-Seq, we show that both types of GO interacted with the plasma membrane and were taken up by the cells in a time- and dose-dependent manner. Both GO types were internalized *via* multiple pathways by BEAS-2B cells; however, macropinocytosis was mainly used for the uptake of s-GO, while CME and CavME were mainly used for the uptake of us-GO. The observed difference was connected with the broad size distribution of the two materials, thoroughly characterized both in water and in the biologically relevant cell culture medium. Furthermore, validation of the uptake pathway was performed by investigating the interference of both types of GO with the uptake of 0.1 and 1 μm beads. We clearly show that s-GO could reduce the uptake of the beads irrespective of their sizes, while us-GO was only capable of reducing the uptake of 0.1 μm beads only. Importantly, we report the shift in the main uptake mechanism from macropinocytosis to clathrin-dependent endocytosis for s-GO at 24 h. This is achieved through activation of mTORC1/2 that negatively regulates macropinocytosis. Finally, both types of GO materials were found to end up in the lysosomes for up to 48 h. This study provides valuable insight into the way GO can be further exploited both biomedically but also in the studies aiming to establish its safety profile.

## Experimental

### Production and characterisation of graphene oxide

Aqueous dispersions of s-GO and us-GO were prepared as described in our previous studies^4,79^ by a modified Hummer’s method coupled with sonication. We used sterilised glassware and GO suspensions were always handled within fume hoods. In brief, 0.8 g of graphite powder (Sigma Aldrich) were mixed with 0.4 g NaNO_3_ and 18.4 mL H_2_SO_4_ (99%) by a 10 minutes rigorous stirring at low temperature (ice bath), followed by the addition of 2.4 g KMnO_4_.The mixture was stirred continuously for 30 minutes (water bath) until a green, homogenous and thick paste was formed. Then, a volume of 37.5 mL water for injections was added dropwise to the reaction volume, while carefully monitoring the temperature rise. The mixture was then stirred for 30 minutes at 98 °C (oil bath). Lastly, the reaction was stopped by the addition of 12.5 mL of H_2_O_2_ and the mixture was left to settle for 1 hour. Subsequently, the dispersion was subjected to a series of washes with water for injections (8800xg, 20 minutes), in order to neutralise the pH, remove the impurities and separate the GO from the graphitic residues. Upon the last two washing steps, GO was separated by vortexing and then solubilised in with warm water for injection (50 °C) from the orange gel layer, which formed at the top of the graphite oxide. Any graphitic residues still present in the dispersion were removed by an additional centrifugation step (8800xg, 20 minutes), at 24 hours post-reaction. The size reduction to small and ultra-small flakes was carried out by sonication (80 W, 45 kHz) for 5 minutes and 4 hours, respectively. The purification was done by centrifugation at 13000 rpm for 5 minutes in the case of s-GO and 1 hour in the case of us-GO.

### UV-visible spectroscopy (UV-vis)

Spectra of GO dilutions in Milli-Q water with concentrations ranging from 2.5 to 20 *μ*g/mL were acquired using a Cary 50 Bio UV-vis spectrophotometer (Varian Inc., Agilent Technologies, UK). Measurements were performed at room temperature in a quartz cuvette (1 mL volume, 1 cm path length). Milli-Q water was used as a blank.

### Fluorescence spectroscopy

Different concentrations of GO dispersions (25–200 *μ*g/mL) were measured with a Cary Eclipse fluorescence spectrophotometer (Varian Inc., Agilent Technologies, UK). Spectra were acquired at room temperature, with *λ*_exc_ set to 525 nm. Milli-Q water was used as a blank.

### Raman spectroscopy

Measurements were recorded by a DXR micro-Raman spectrometer (Thermo Fisher Scientific, UK) equipped with a 633 nm laser set to 0.4 mW. Calibration was done on a polystyrene standard, the chosen objective was 50x, and the pinhole was set to 50 μm. Spectra were then recorded between 500 and 4000 cm^−1^ with a resolution of 2.5 cm^−1^. All spectra were processed by background subtraction and normalisation by the G band intensity using OriginPro 8.5.1 software.

### Zeta Potential measurements

The Zeta potential values of GO suspensions in Milli-Q water were measured with a ZetaSizer Nano ZS instrument (Malvern, UK). All measurements were performed at 25 °C using disposable folded capillary zeta cells. The results are reported as the average ± standard deviation of three measurements per sample.

### Thermogravimetric analysis (TGA)

The oxidation degree of GO materials was extracted from the degradation patterns measured with a TGA 4000 thermogravimetric analyser (PerkinElmer Ltd, UK). All measurements were done on 2 mg lyophilised material, in a Nitrogen atmosphere (20 mL/min) and at temperatures ranging from 25 to 995 °C (10 °C/min). The material residues remained at 995 °C were burned by switching the purge gas to Oxygen, for 15 minutes.

### X-ray photoelectron spectroscopy (XPS)

XPS measurements of lyophilised GO samples were analysed using a Thermo Theta Probe XPS spectrometer with a monochromatic Al K-*α* source of 1486.68 eV. The spectra were acquired with PE of 40 kV, 0.1 eV step size and an average of 20 scans. Spectra post-processing was done with CasaXPS software (Casa Software Ltd, UK). A Shirley background subtraction was applied to all spectra and Gaussian–Lorentzian (70:30) functions were used for fitting the functional groups, except for the asymmetric C–C and C=C peak, which was fitted using an asymmetric Lorentzian function. The full width half maximum (FWHM) value was constrained between 0.5 and 2 eV for all peaks, except for the *π–π*\*. The following constrain regions were set for the binding energies: 284-285.5 for C-C/C=C, 285.5-286.5 for C-O, 286.8-287.8 for C=O, 288.6-290 for COOH and >290 for π-π*.

### Transmission electron microscopy (TEM)

TEM analysis was performed on an FEI Tecnai 12 BioTwin equipment (FEI, Eindhoven, NL) with the electron beam set to 100 kV. The samples were prepared on 300-mesh carbon-coated copper grids, at room temperature and in a clean environment. A volume of 20 μL of GO dispersion was drop-casted on the grid, and the excess was removed after 1 minute with filter paper, leaving a thin layer of suspension to fully dry. Eventually, a small drop was casted and left to dry at the edge of the grid. Images were captured with an AMT digital camera (Gatan, UK). The raw data were further processed using ImageJ the lateral size of the GO flakes was manually measured by determining the longest Feret diameter in each flake.

### Atomic force microscopy (AFM)

A multimode atomic force microscope (Bruker, UK) was used in tapping mode, using Otespa-R3 probes (Bruker, UK). Samples were prepared on poly-L-lysine 0.01% (Sigma Aldrich P4707) coated mica substrates, by drop-casting a volume of 20 μL of 100 μg/mL GO dilution in Milli-Q water for 1 minute, followed by a washing step with 1 mL Milli-Q water and drying overnight in a drying cabinet (37°C). Scanning parameters were set as follows: 1 Hz scanning rate, 250 mV amplitude set-point, 512 lines per scan, an integral gain of 1 and a proportional gain of 5. Images were processed with the Bruker Nanoscope Analysis software-Version 1.4; the lateral size of the GO flakes was manually measured by determining the longest Feret diameter in each flake.

### Characterisation of GO in cell culture medium

GO (50 μg/mL) was incubated in RPMI-1640 cell culture medium (R8758, Sigma-Aldrich, Merck Sigma, UK), supplemented with 10% FBS (F9665, Sigma-Aldrich, Merck Sigma, UK), 1000 units Penicillin, and 1 mg/mL Streptomycin (Sigma-Aldrich, Merck Sigma, UK) for a series of time points (0 min, 4 h and 24 h). At indicated time point, GO was centrifuged (30 min, 13 000 rpm), suspended in Milli-Q water (1 mL), re-centrifuged (30 min, 13 000 rpm), and then re-suspended in Milli-Q water (1 mL) for Zeta-potential and size analysis using ZetaSizer Nano ZS instrument (Malvern, UK).

### Cell culture

Human epithelial bronchial immortalized cells (BEAS-2B, CRL-9609, ATCC, LGC standards, UK) were maintained in RPMI-1640 cell culture medium, mouse fibroblast embryonic immortalized cells (NIH/3T3, CRL-1658, ATCC, LGC standards, UK), human epithelial keratinocyte immortalized cells (HaCaT, PCS-200-011, ATCC, LGC standards, UK) and human epithelial embryonic kidney immortalized cells (293T, CRL-2316, ATCC, LGC standards, UK) were maintained in DMEM cell culture medium (D6429, Sigma-Aldrich, Merck Sigma, UK), all supplemented with 10% FBS, 1000 units Penicillin, and 1 mg/mL Streptomycin at 37 °C in a humidified 5% CO_2_ incubator. Cells were split at 80% confluence with 0.05% Trypsin-EDTA (Sigma-Aldrich, Merck Sigma, UK), and 10% FBS was used to stop the activity of Trypsin-EDTA.

### Cell culture treatments

Depending on the experiment cells were seeded in either Cellview™ dishes (for all confocal microscopy related experiments) or in 12-well plates (for all flow cytometry related experiments, and GO deposition measurement). Cells were treated at 60-80% confluence unless stated otherwise. Cells were always seeded in the cell-type specific growth medium up to 24 h before treatments/pre-treatments, and in RPMI-1640 cell culture medium for all treatments/pre-treatments, all supplemented with 10% FBS, 1000 units Penicillin, and 1 mg/mL Streptomycin at 37 °C in a humidified 5% CO_2_ incubator.

### Evaluation of GO deposition on cells

BEAS-2B cells were seeded in 12 well plates and treated with s-GO or us-GO for 4 or 24 h (0, 25 and 50 μg/mL, 1mL/well) in RPMI medium free of phenol red supplemented with 10 % of FBS (Gibco). After 4 and 24 hours, supernatants were collected and absorbance values were measured using UV-VIS spectrometry (Cary 50 Bio, Varian). For each independent experiment and nanomaterial, standard curves were prepared with concentrations ranging from 0 to 50 ug/mL and concentrations of material remaining in supernatants were determined. Absorbance values were measured at 750 nm to avoid interferences from the culture medium. Mass deposited were calculated by subtracting the mass remaining in the supernatant from the mass administered. Percentages of deposition were also calculated. Potential differences of deposition between s-GO and us-GO were evaluated using one-way ANOVA followed by Dunnett’s post hoc test (minimum of 3 independent experiments).

### Confocal microscopy

#### Uptake of GO

Cells were treated with s-GO or us-GO (25, 50 and 75 μg/mL, 0.5 mL/well) for 24 h. After 24 h of treatment, supernatants were removed and replaced by CellMask™ green plasma membrane stain (C37608, Thermo Fisher Scientific, UK) prepared in the control medium (dilution 1:2500). Cells were then examined using Zeiss 780 CLSM using the 40X objective. Images were then processed using Zeiss microscope software ZEN. Excitation/emission wavelength: CellMask™ green = 488/520, GO = 594/620-690 nm. Quantitative characterization for the uptake of GO in cells was carried out using the ImageJ software. Cells from 3 independent set of confocal images were analysed and standardised. Statistical analysis of the result was performed using GraphPad Prism (version 8) with analysis of variance (one-way ANOVA) and Tukey’s multiple comparisons test. Differences with p < 0.05 were considered as statistically significant: **** p < 0.0001.

#### Uptake of 1μm beads in the presence or absence of GO

BEAS-2B cells were pre-treated at ~30-40% confluence with either complete RPMI-1640 cell culture medium (0.5 mL/well) or s-GO/us-GO (50 μg/mL, 0.5 mL/well) and incubated for 4 h. After pre-treatments, 1 μm beads (1.5 μL/mL, 0.5 mL/well, F8814, Thermo Scientific, UK) or complete RPMI-1640 cell culture medium (0.5 mL/well) containing CellMask™ green plasma membrane stain (dilution 1:2500) were added to the cells and incubated for another 24 h. Cells were then examined using Zeiss 710 CLSM (40X objective, imaging mode) and processed using Zeiss microscope software ZEN. Excitation/emission wavelength: CellMask™ green = 488/520, 1 μm beads = 365/415.

#### Subcellular localization of GO

BEAS-2B cells were incubated with CellLight™ Lysosomes-GFP, BacMam 2.0 at 40% confluence (20 μl of CellLight™ Lysosomes-GFP was diluted in 0.5 ml of complete cell culture medium/well; C10596, Thermo Fischer, UK) for 16 h (overnight). After incubation, cells were washed once with PBS (with Ca^2+^/Mg^2+^) and treated with 50 μg/mL of s-GO or us-GO for 24h. After the treatment cells were imaged using Zeiss 780 live-cell time lapse confocal microscope for the interval of 20 min. Excitation/emission wavelength for CellLight™ Lysosomes-GFP = 488/520.

#### Treatments with pharmacological inhibitors

BEAS-2B cells were pre-treated at ~80-90% confluence with EIPA (100 μM, 1 mL/well, A3085, Sigma-Aldrich, Merck Sigma, UK), monodansylcadavarine (55 μM, 1 mL/well, D4008, Sigma-Aldrich, Merck Sigma, UK), chlorpromazine (20 μM, 1 mL/well, *C8138*, Sigma-Aldrich, Merck Sigma, UK), genistein (350 μM, 1 mL/well, G6649, Sigma-Aldrich, Merck Sigma, UK), cytochalasin D (0.5 μM, 1 mL/well, C8273, Sigma-Aldrich, Merck Sigma, UK), latrunculin A (100 nM, 1 mL/well, L5163, Sigma-Aldrich, Merck Sigma, UK), sodium azide (100 mM, 1 mL/well, 26626-22-8, Sigma-Aldrich, Merck Sigma, UK) or dynasore (300 μM, 1 mL/well, *D7693*, Sigma-Aldrich, Merck Sigma, UK) for 30 min. After pre-treatment, supernatants were removed and cells were treated with s-GO (50 μg/mL, 1 mL/well) or us-GO (50 μg/mL, 1 mL/well) prepared in the corresponding pre-treatment solution containing inhibitor and incubated for 4 h. Cells were then detached with 0.05% Trypsin-EDTA (300 μL/well, 10 min,), neutralised with 10% FBS (30 μL/well), collected in 1.5 mL tube, stored in ice and analysed by FACSVerse flow cytometry using the PE-Cy7-A channel (bandpass: 488 780/60). Excitation/emission wavelength: GO = 594/620-690.

#### Staining of actin filaments

BEAS-2B cells were treated with the EIPA (100 μM, 0.5 mL/well), monodansylcadavarine (55 μM, 0.5 mL/well), chlorpromazine (20 μM, 0.5 mL/well), genistein (350 μM, 0.5 mL/well), cytochalasin D (0.5 μM, 0.5 mL/well), latrunculin A (100 nM, 0.5 mL/well), sodium azide (100 mM, 0.5 mL/well) or dynasore (300 μM, 0.5 mL/well) for 4 h and 30 min. After treatment, cells were washed two times with pre-warmed PBS (0.5 mL/well, D8662, Sigma-Aldrich, Merck Sigma, UK), fixed with formaldehyde (3.7%, 0.5 ml/well, 10 min; 28908 Thermo Fisher Scientific, UK). After fixation cells were permeabilized with Triton-X (0.1 % in PBS, 0.5 mL/well, 5 mins) and washed two times with PBS (0.5 mL/well)before staining with Alexa Fluor™ 488 Phalloidin (A12379, Thermo Fischer, UK) prepared in PBS (in a dilution of 1:1500) for 20 min. Cells were washed two times with PBS (0.5 mL/well), following by the addition of ProLong™ Gold Antifade Mountant (P36930, Thermo Fisher Scientific, UK) and covered with a coverslip. Cells were then observed using Zeiss 780 CLSM using the 40X objective. Images were processed using Zeiss microscope software ZEN. Excitation/emission wavelength: Phalloidin = 495/518.

### Flow cytometry

#### Cellular interactions with GO

BEAS-2B cells were treated with s-GO or us-GO (25, 50 and 75 μg/mL, 1mL/well) for 4 h or 24 h. Cells were then detached with 0.05% Trypsin-EDTA (300 μL/well, 10 min), neutralised with 10% FBS (30 μL/well), collected in 1.5 mL tube, stored in ice and analysed by FACSVerse flow cytometry using the PE-Cy7-A channel (band pass: 488 780/60). Excitation/emission band pass: GO = 594/620-690.

#### Cellular interactions with the beads in the presence or absence of GO

BEAS-2B cells were pre-treated with either complete RPMI-1640 cell culture medium (1 mL/well) or s-GO/us-GO (50 μg/mL, 1 mL/well) at ~30-40% confluence for 4 h. After pre-treatments, 0.1 μm beads (1.5 μL/mL, 1 mL/well, F8803, Thermo Scientific, UK), 1 μm beads (1.5 μL/mL, 1 mL/well, F8823, Thermo Scientific, UK) or complete RPMI-1640 cell culture medium (1 mL/well) were added to the cells and incubated for another 24 h. Cells were then detached with 0.05% Trypsin-EDTA (300 μL/well, 10 min), neutralised with 10% FBS (30 μL/well), collected in 1.5 mL tube, stored in ice and analysed by FACSVerse flow cytometry (bandpass: 488 530/30) or Fortessa X20 (bandpass: 488 529/24) using the FITC channel. Trypan Blue (0.2 % solution) was added to each sample just before analysis. Excitation/emission wavelengths: 0.1 μm beads = 505/515, 1 μm beads = 505/515.

#### PI/AV assay

BEAS-2B cells were treated at ~80% confluence with EIPA (100 μM, 1 mL/well,), monodansylcadavarine(55 μM, 1 mL/well,), chlorpromazine (20 μM, 1 mL/well, *C8138*), genistein (350 μM, 1 mL/well), cytochalasin D (0.5 μM, 1 mL/well), latrunculin A (100 nM, 1 mL/well), sodium azide (100 mM, 1 mL/well) or dynasore (300 μM, 1 mL/well) for 4 h and 30 min. Cells were then detached with 0.05% Trypsin-EDTA (300 μL/well, 10 min), neutralised with 10% FBS (30 μL/well) and collected in 1.5 mL tube. Cells were centrifuged (1500 rpm, 5 min) and re-suspended in diluted Annexin-binding buffer (1X, 200 μL/tube, V13246, Thermo Fisher Scientific, UK). Cells were labelled with Annexin V (1 μL/tube, A13201, Thermo Fisher Scientific, UK) for 20 min. Samples were stored on ice and analysed by FACSVerse flow cytometry using the FITC-A (bandpass: 488 530/30)and PE-A (bandpass: 488 574/26) channel, Propidium Iodide (1 μL/tube, P4864-10ML, Sigma-Aldrich, Merck Sigma, UK) was added to the cells just before analysis of the samples. Excitation/emission wavelength: Annexin V = 495/519, Propidium Iodide = 493/636.

#### TEM

BEAS-2B cells were grown on a sterilized Aclar placed in a 12-well plate and treated with 50 μg/mL of s-GO or us-GO when they reached 70% confluence. After 24 h of treatment, cells were fixed at room temperature using 4% glutaraldehyde/4% paraformaldehyde prepared in 0.2 M HEPES buffer for at least 2 h, washed three times using ddH_2_O, then incubated for 2 h in ferrocyanide reduced osmium. After dehydration in increasing concentrations of ethanol (from 30% up to 100%) and then two times in acetone (100%, 30 min) the samples were immersed in an increasing concentration of TAAB 812 resin in acetone. Ultrathin sections of 80 nm, obtained with a diamond knife using a Leica U6 ultramicrotome, were mounted on the grids and before being examined with an FEI TECNAI transmission electron microscope.

#### RNA-Seq analysis

BEAS-2B cells were treated with s-GO (50 μg/mL for 24 h). RNA was extracted using a PureLink RNA mini kit (Invitrogen). RNA-Seq libraries were generated using the TruSeq Stranded mRNA assay (Illumina, Inc.) according to the manufacturer’s instructions. Briefly, poly-T, oligo-attached, magnetic beads were used to extract polyadenylated mRNA from 1 μg of total RNA. The mRNA was then fragmented using divalent cations under high temperature and then transcribed into first strand cDNA using random primers. Second strand cDNA was then synthesized using DNA polymerase I and RNase H, and a single “A” base addition was performed. Adapters were then ligated to the cDNA fragments and then purified and enriched by PCR to create the final cDNA library. Adapter indices were used to multiplex libraries, which were pooled prior to cluster generation using a cBot instrument. The loaded flow cell was then pair-end sequenced (101 + 101 cycles, plus indices) on an Illumina HiSeq4000 instrument. Demultiplexing of the output data (allowing one mismatch) and BCL-to-Fastq conversion was performed with CASAVA 1.8.3. Sequencing quality for each sample was determined using the FastQC program. Low-quality sequence data were removed utilizing the trimmomatic program. STAR v2.4.0 was utilized to map the trimmed sequence into the human genome. Raw counts for each sample were generated by the htseq-count program and subsequently normalized relative to respective library sizes using DESeq2 package for the R statistical program^80,81^. The DESeq2 program was additionally used to plot the PCA with all sample data to visualize different clusters at multiple levels that describes the maximum variance within the data set. Genes of interest were identified by pairwise comparisons. False discovery rate (FDR) adjusted p values were used to evaluate significance.

#### Functional and Pathway Enrichment Analysis

Genes with an FDR-corrected p value of less than 0.05 used for enrichment analyses exploring GO Term Pathway and Panther databases utilizing the Enrichr gene set enrichment analysis web Server^82^. Gene lists of significant features (*p* < 0.05) were then confirmed through literature search to be integral to the pathways identified and further investigated for consistency in response direction to identify the key mechanism involved in the s-GO response. Gene lists found to be enriched in these features were then used to generate heat maps utilising pheatmap^83^ R-package^81^ to visualize the effects of s-GO.

#### Statistical analysis

All experiments were repeated at least two times with duplicates, triplicates or quintuplicates for each condition, and the results were expressed as mean ± standard deviation. Flow cytometry data were analysed using GraphPad Prism (version 7) with analysis of variance (two-way ANOVA) and *post hoc* Sidak’s multiple comparisons test. Differences with *p* < 0.05 were considered as statistically significant: * *p* < 0.05, **** *p* < 0.0001. Deposition of s-GO and us-GO on BEAS-2B cells were evaluated using one-way ANOVA followed by Dunnett’s post hoc test.

## Acknowledgements

Authors would like to acknowledge the funding from Graphene Flagship WP4 project (FP7-ICT-2013-FET-F-604391) and from Horizon 2020 research and innovation programme under the grant agreement no. 785219 (Graphene Flagship Core2). Both Y.C. and L.E.C would like to acknowledge the studentship from the Engineering and Physical Sciences Research Council (EPSRC) Centre for Doctoral Training programme (Graphene NOWNANO CDT; EP/L01548X/1). The authors acknowledge the staff of the Faculty of Biology Medicine and Health EM Facility, for their expertise and assistance, and the Wellcome Trust for equipment grant support to the EM Facility. The University of Manchester Bioimaging and Single Cell Genomics Facility microscopes used in this study were purchased with grants from the Biotechnology and Biological Sciences Research Council (BBSRC), Wellcome Trust, and the University of Manchester Strategic Fund. The authors also wish to thank Dr. N. Hodson from the Bio-AFM Facility for assistance and advice regarding the AFM instrumentation. The authors acknowledge the Manchester Collaborative Centre for Inflammation Research (MCCIR) as the funding source for the FACSVerse and Fortessa X20 instrument. Authors also acknowledge EPSRC Harwell XPS facility based at Cardiff University for the provision of X-ray photoelectron spectra.

## Supporting Information

**Figure S1.**
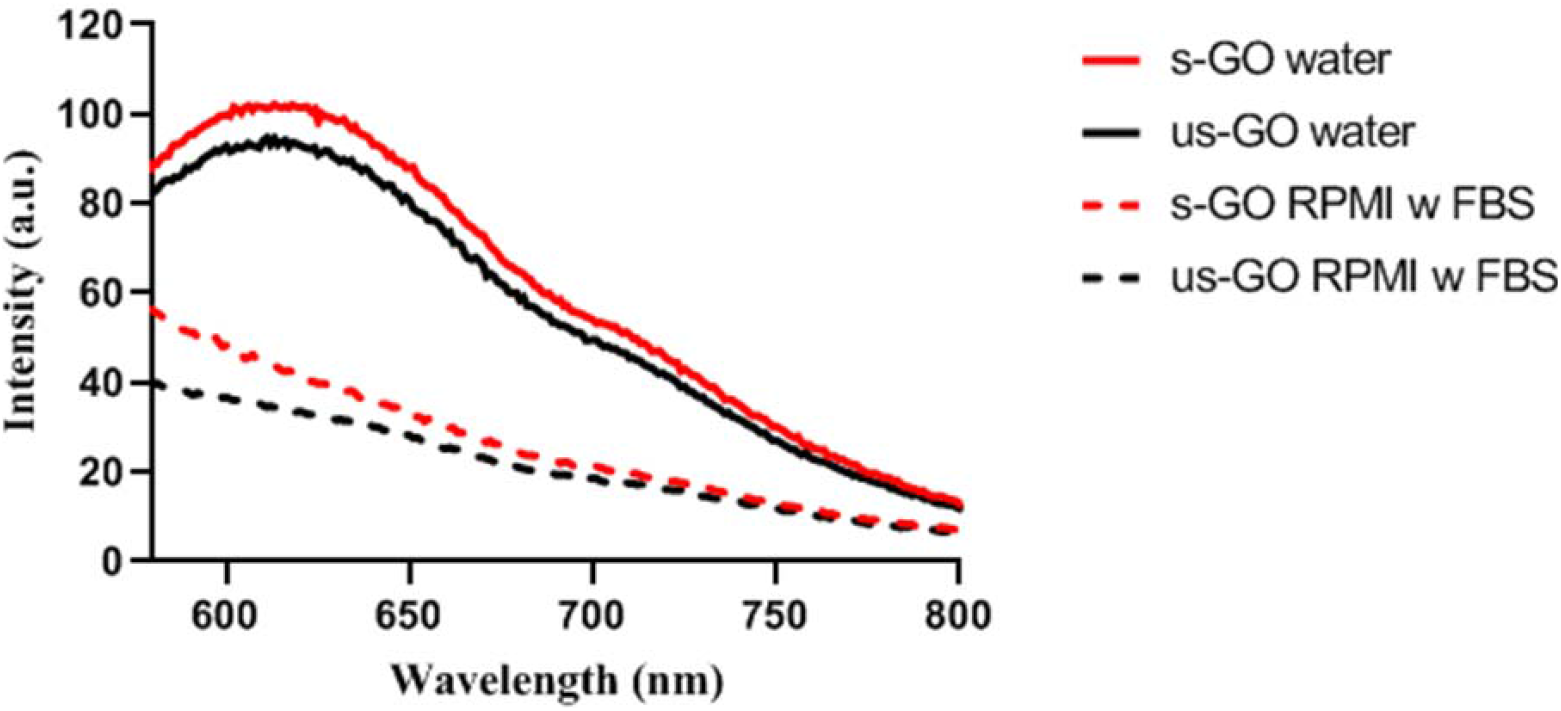
Spectrofluorometric analysis of the emission spectra of s-GO and us-GO (2 mg/mL) dispersed in water (solid lines) and in RPMI with 10 % serum (patterned lines – samples prepared in RPMI with serum were re-suspended in water for the measurement) using an excitation wavelength of 525 nm. This result shows that s-GO retained a higher intensity of intrinsic fluorescence comparing to us-GO in both water and cell culture medium with serum. The lower fluorescence intensity of the material in the cell culture medium can be explained by the agglomeration of material in the culture medium.

**Figure S2.**
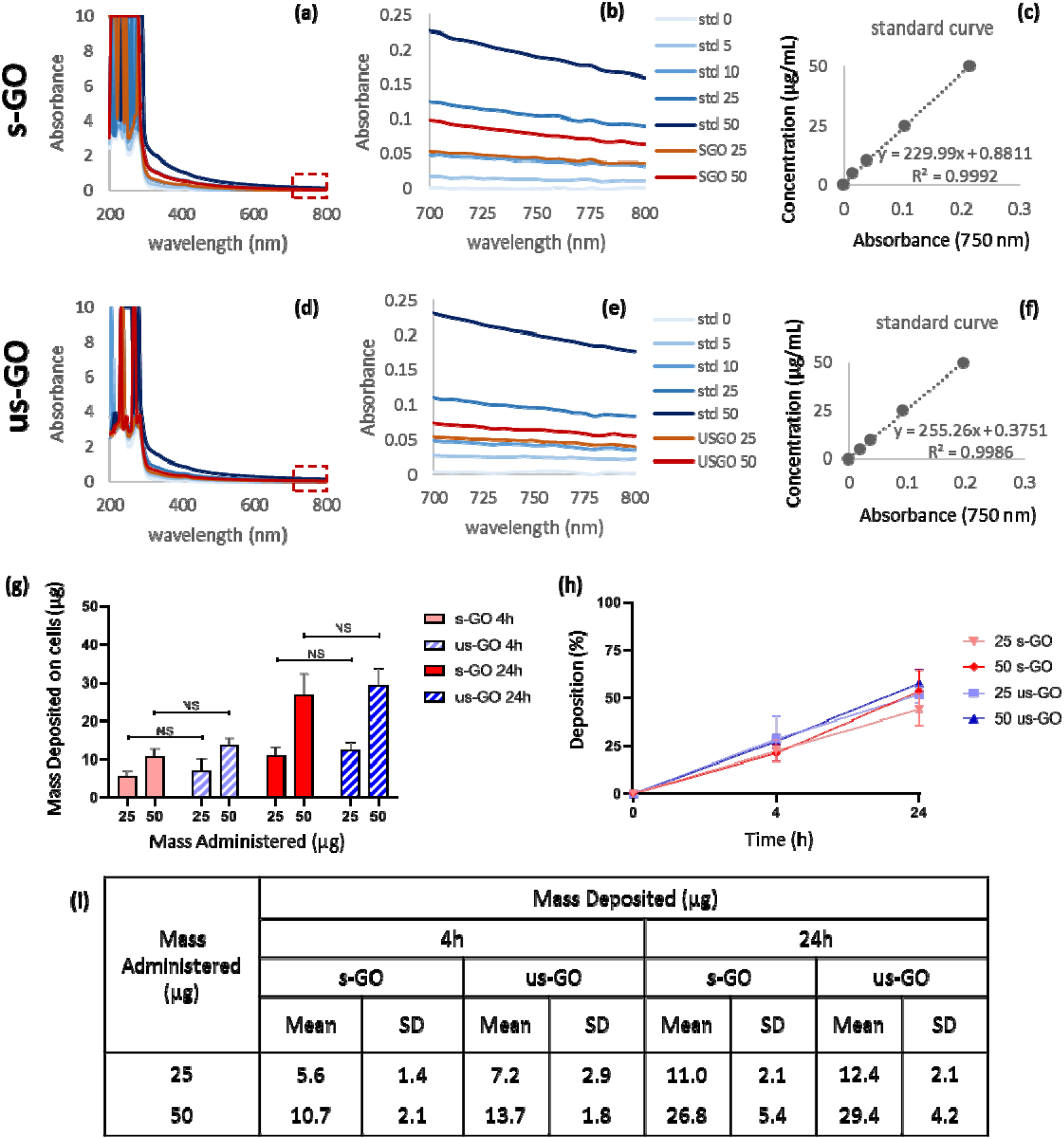
GO deposition measurement on BEAS-2B cells. To evaluate the mass of s-GO and us-GO deposited on BEAS-2B cells, supernatants were collected at 4 and 24 hours and the absorbance was measured using UV-VIS absorption spectroscopy. UV-vis spectra of (**a** and **b**) s-GO and (**d** and **e**) us-GO at concentrations ranging from 0 to 50 μg/mL is showed, with enlarged spectra at region of interest (**b** and **e**). The concentrations of materials remaining in suspension were determined thanks to standard curves (**c** and **f**), based on the absorbance values measured at 750 nm. The mass of material deposited on cells (**g**) were then calculated based on the difference between the mass of material administrated and remaining in suspension (*Mass Deposited = Mass Administrated – Mass remained in suspension*). Percentages of deposition (**h**) were also calculated. Difference of deposition between s-GO and us-GO were evaluated using one-way ANOVA followed by Dunnett’s post hoc test (minimum of 3 independent experiments). No significant differences in mass deposited were observed between the s-GO and us-GO. (*std = standard, SD = standard deviation, NS = not statistically significant*)

**Figure S3.**
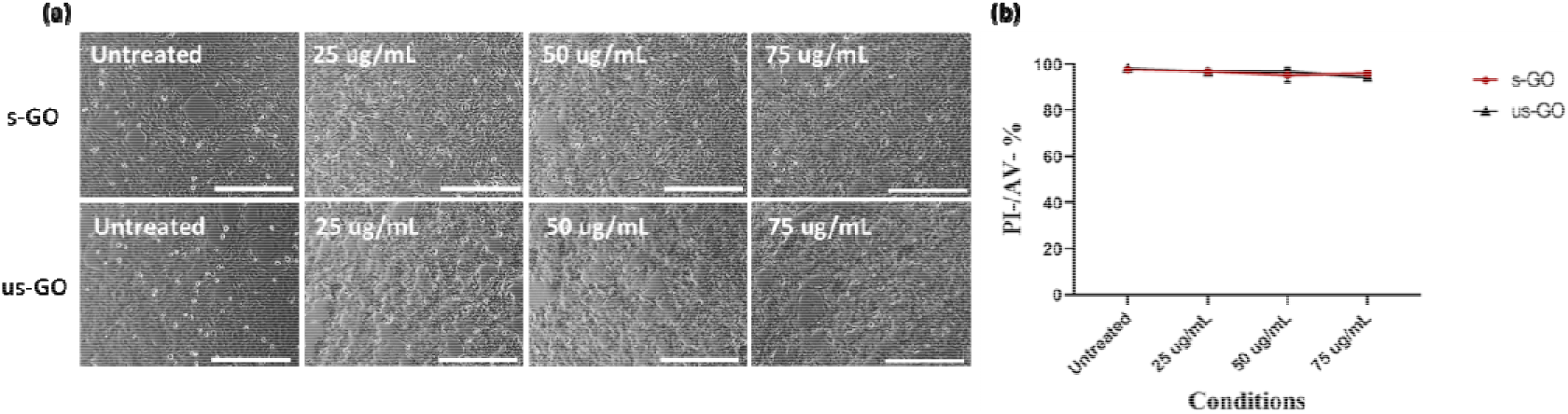
Cytotoxicity of s- and us-GO in BEAS-2B (24 h treatment) was assessed *via* (**a**) optical imaging, and (**b**) propidium iodide (PI)/ annexin V (AV) staining using flow cytometry (positive control is shown in **Figure S10**). Representative optical images of BEAS-2B cells treated with and without s- or us-GO are showed. And we observed no obvious changes in cell morphology and cell confluence with the treatment of GO compared to the untreated cells. The PI/AV assay confirmed no significant difference in the percentage of live cells (i.e. cells unstained by both PI/AV (PI-/AV-)) with the treatment of GO compared to the untreated cells. Scale bar = 400 μm.

**Figure S4.**
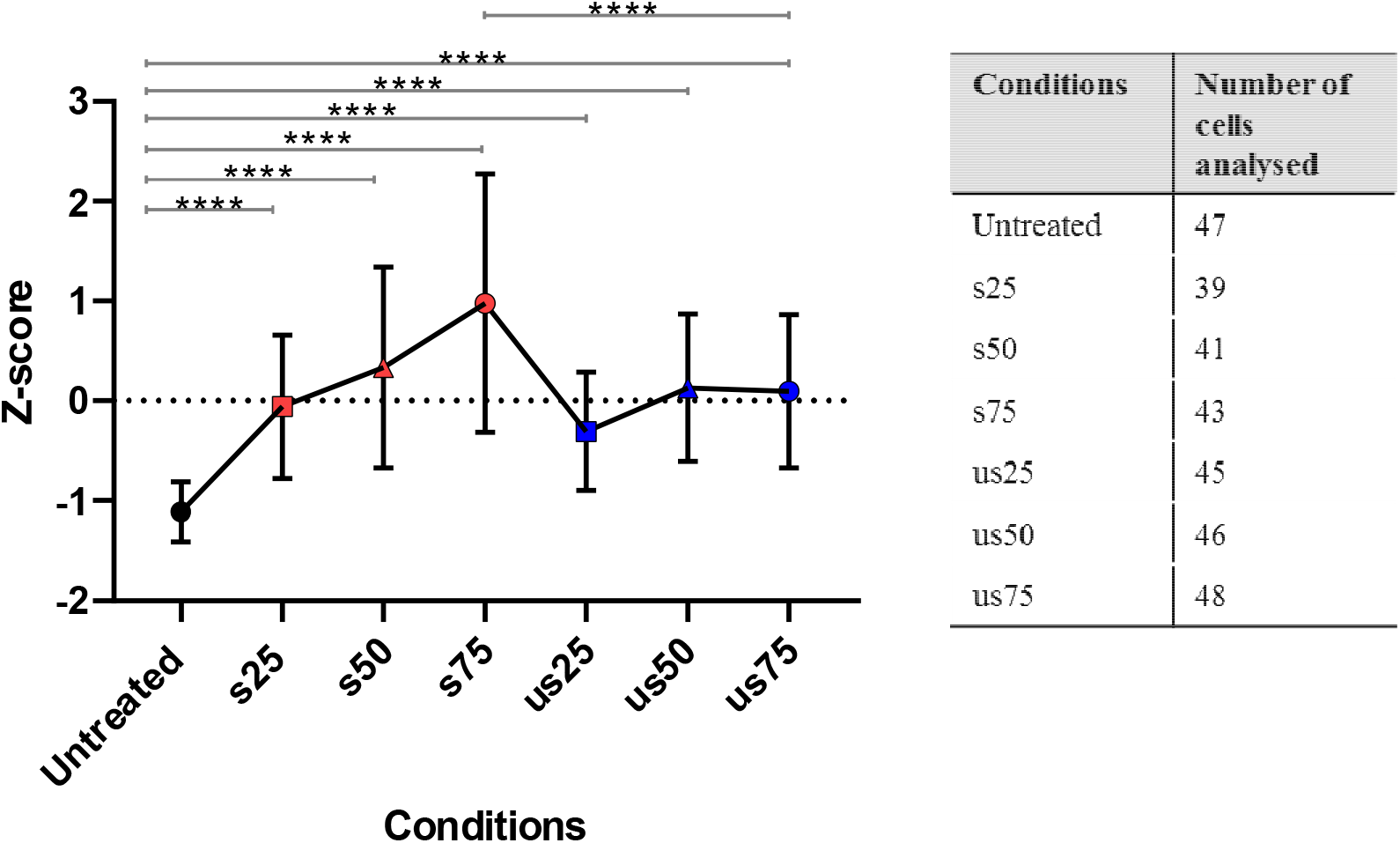
Standardised quantitative analysis of the fluorescent images for the interaction of GO with BEAS-2B cells. The results showed the uptake of GO in treated samples (regardless of size and concentrations) are all significantly greater than untreated samples. And the result showed no statistical differences in the uptake of s- and us-GO except for the highest concentration of 75 μg/mL. The images were manually analysed using ImageJ. The data were statistically analysed using analysis of variance (one-way ANOVA) and Tukey’s multiple comparison test with **** equate to p < 0.0001. n = 3 independent experiments. s = s-GO, us = us-GO, 25 = 25 μg/mL, 50 = 50 μg/mL and 75 = 75 μg/mL.

**Figure S5.**
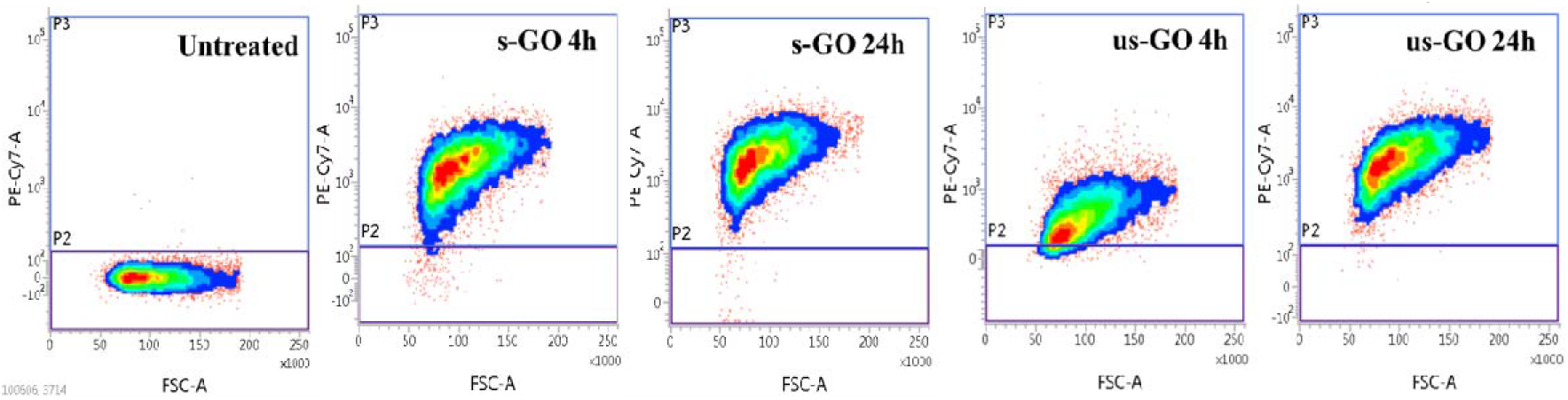
FACS density plots of untreated and BEAS-2B cells treated with 75 μg/mL of s-GO or us-GO at 4 h and 24 h time points. PE-Cy7-A channel was used to detect the auto-fluorescent signal from the GO. Cellular interaction with GO was assessed by flow cytometry by measuring the auto-fluorescent signal of GO in the PE-Cy7-A channel. It is obvious that the intensity of fluorescence of us-GO is much lower at 4 h compared to 24 h, whereas for s-GO the intensity of fluorescence is only slightly lower at 4 h compared to 24 h.

**Figure S6.**
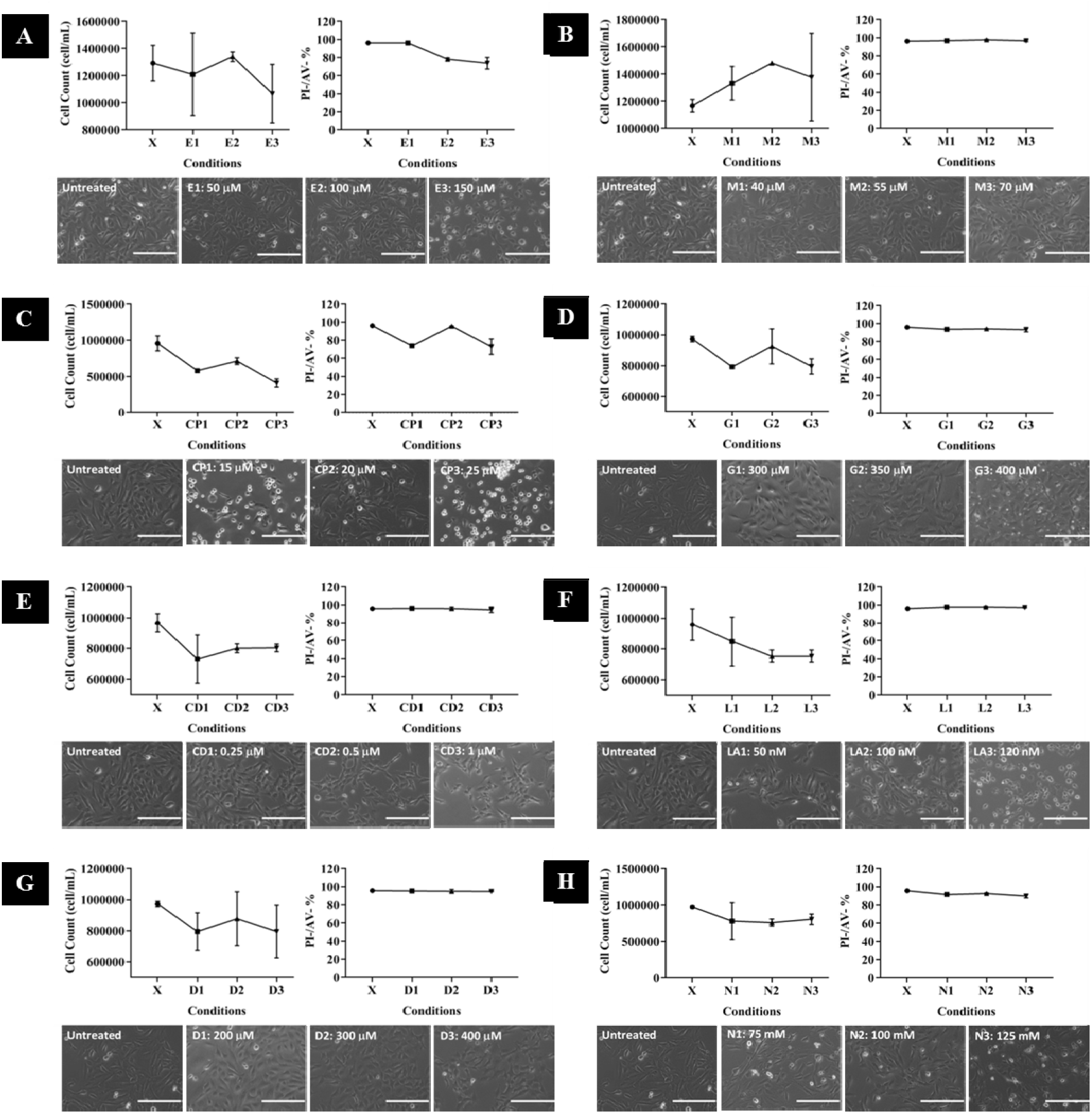
Cytotoxicity of each inhibitor using three different concentrations was assessed *via* optical imaging, cell counting by Trypan Blue dye exclusion assay, and propidium iodide (PI)/ annexin V (AV) staining using flow cytometry (positive control is shown in **Figure S10**). The optical images enabled to assess changes in cell morphology and healthiness of the monolayers, while live cell counting and PI/AV bivariate plots quantified cytotoxicity induced by the treatment with inhibitors. PI will stain for early necrotic cells, whereas AV will stain for early aprotic cells, cells which stained by both PI and AV indicate late cell death and cells unstained by both PI/AV are live cells (PI-/AV-). The selected working concentration for each inhibitor is further tested for disruption to actin filaments (**Figure S7**). Scale bar = 200 μm. (**A**: Ethyl-isopropyl amiloride, **B**: Monodansylcadaverine, **C**: Chlorpromazine, **D**: Genistein, **E**: Cytochalasin D, **F**: Latrunculin A, **G**: Dynasore and **H**: Sodium azide)

**Figure S7.**
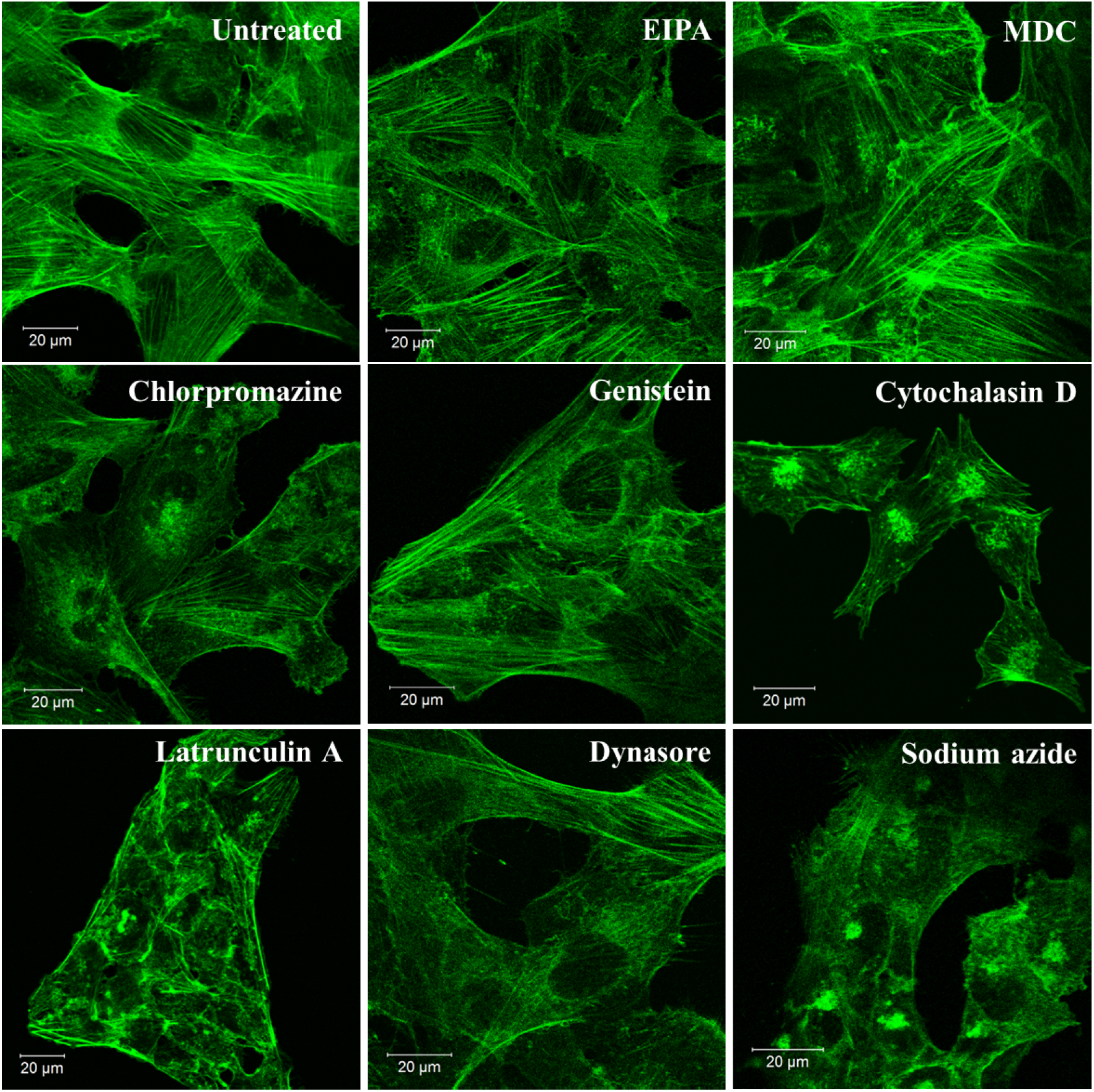
Actin filament staining of BEAS-2B cells with inhibitor at the selected working concentration (**Table 2**) using confocal microscopy. The result shows that in general the inhibitors caused no disruption of actin filaments, except for Cytochalasin D and Latrunculin A. Green = actin filaments.

**Figure S8.**
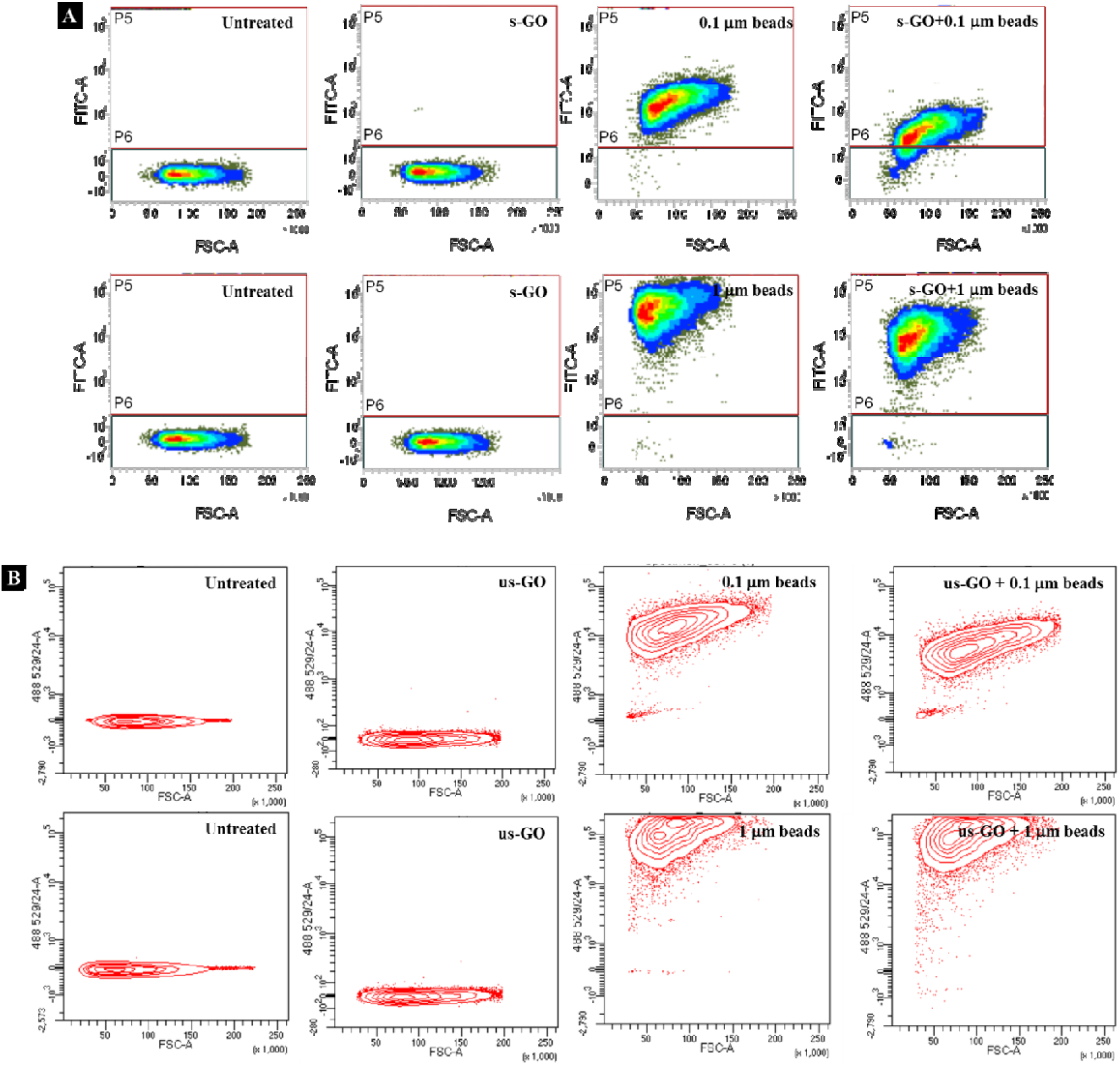
FACS density plot of untreated BEAS-2B cells, BEAS-2B cells treated with only GO, beads (0.1 or 1 μm beads), or beads (0.1 or 1 μm beads) in the presence of (**A**) s-GO, acquired using FACSVerse or (**B**) us-GO, acquired using Fortessa X20 at 50 μg/mL. This figure confirms that the fluorescent signal of the 0.1 and 1 μm beads changes in the presence of s-GO and us-GO, but to a different extent. Fluorescent signal of GO only was comparable to untreated cells.

**Figure S9.**
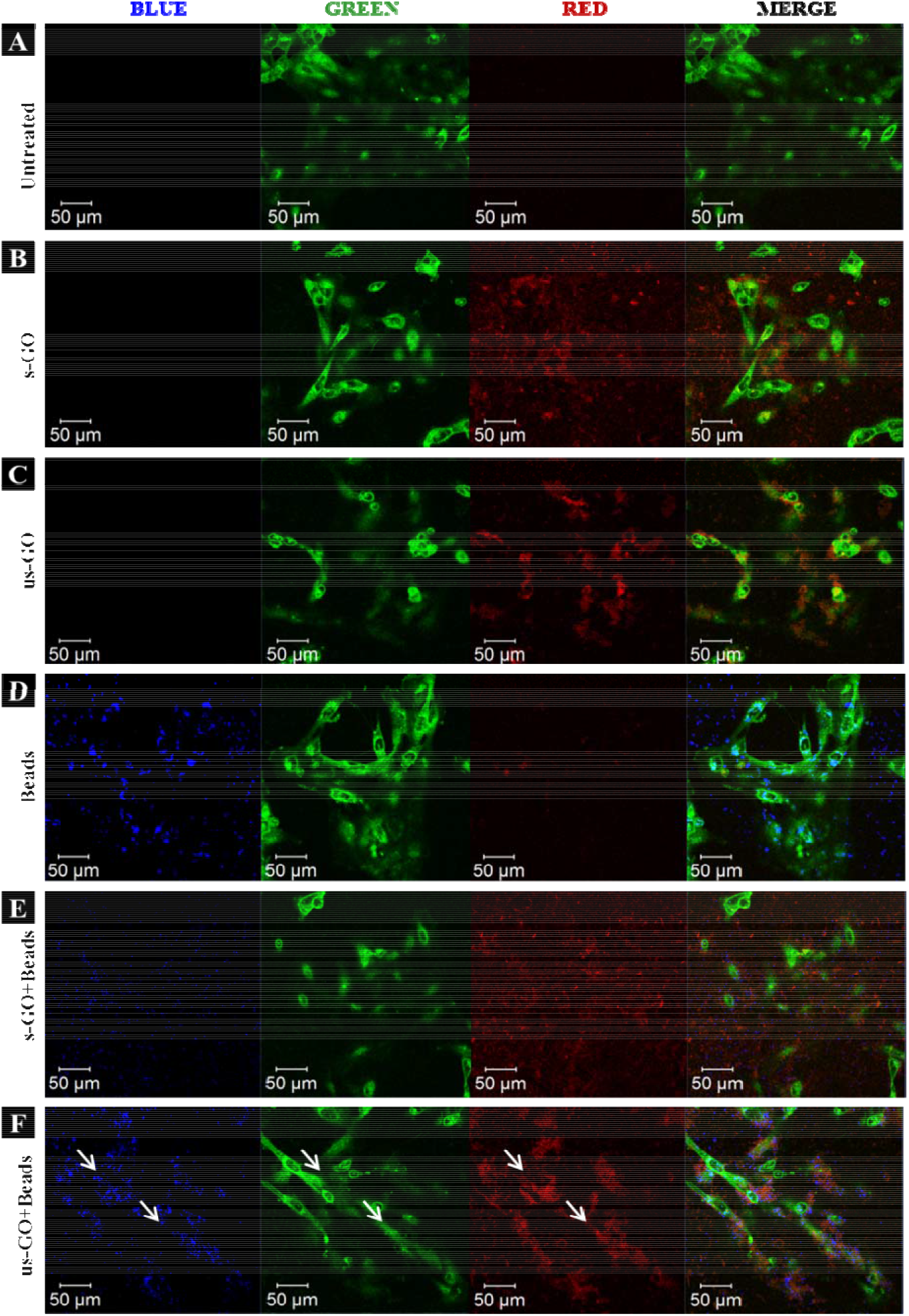
The apical section of BEAS-2B cells treated with 1 μm beads in the presence/absence of s-GO/us-GO (50 μg/mL). White arrows indicate regions co-localisation between the beads and us-GO on top of the cells (**F**), which was not observed for cells treated with s-GO + beads (**E**). (**A**: untreated cells, **B**: s-GO treated, **C**: us-GO treated, **D**: beads treatment, **E**: s-GO + beads, **F**: us-GO + beads) Green = plasma membrane, red = GO, blue = beads.

**Table S1.**
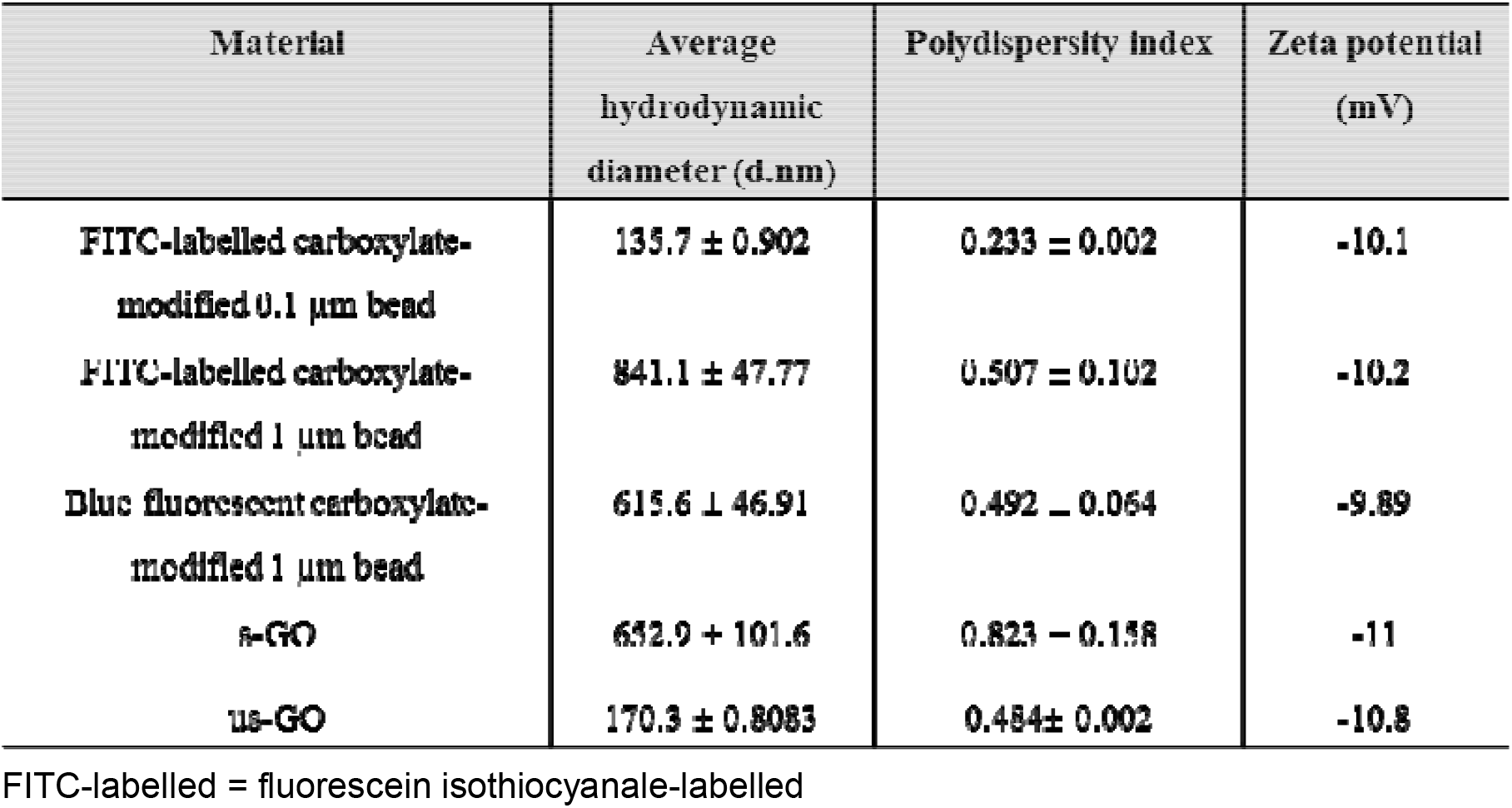
Summary of the sizes and Zeta potentials of the materials (in RPMI w FBS) used in the study. Materials were prepared in RPMI w FBS and measured within 1 hour of sample preparations. The result indicated the 0.1 and 1 μm beads had a similar surface charge to the GO we produced.

**Figure S10.**
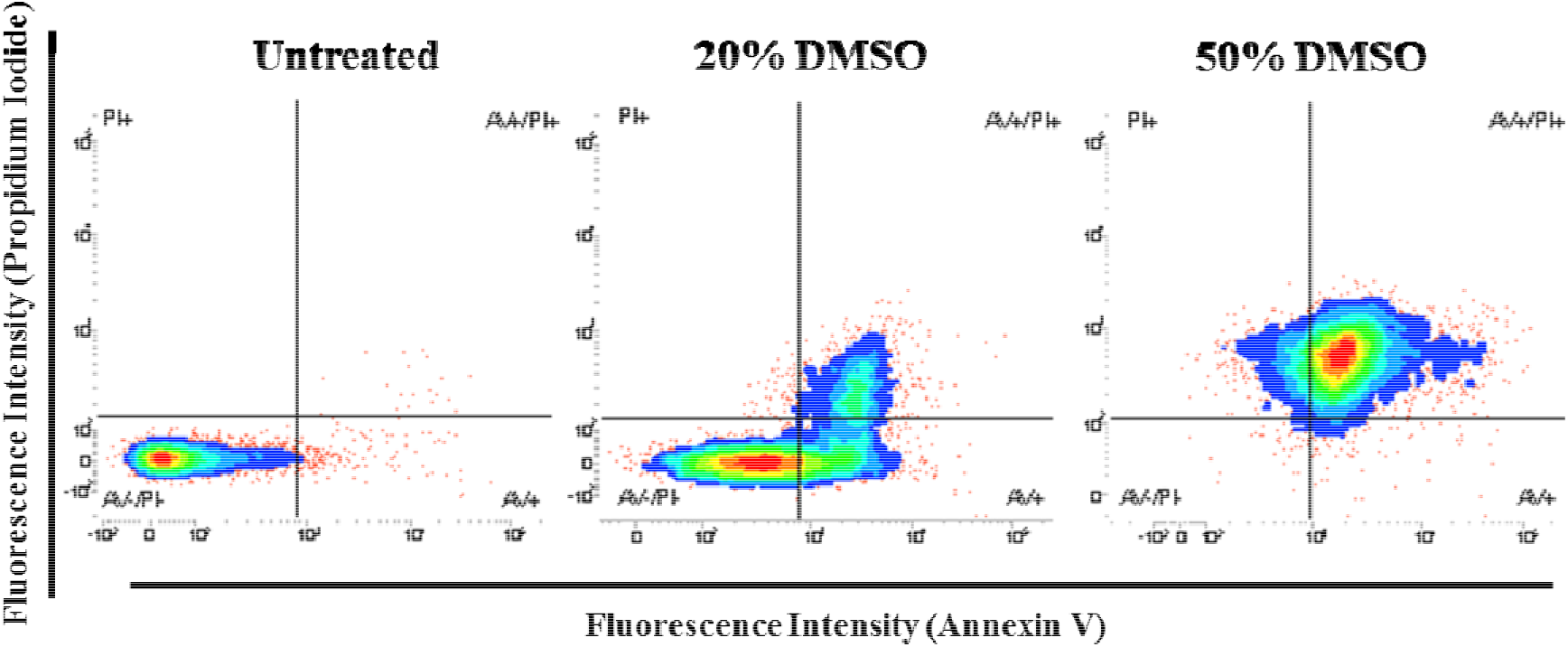
Dimethyl sulfoxide (DMSO) is used as the positive control in propidium iodide (PI)/ annexin V (AV) staining of BEAS-2B cells.

## References

1. D. A. Jasim, N. Lozano, C. Bussy, I. Barbolina, A. F. Rodrigues, K. S. Novoselov and K. Kostarelos, Graphene-based papers as substrates for cell growth: Characterisation and impact on mammalian cells, FlatChem, 2018, 12, 17–25.

2. M. Vincent, I. De Lázaro and K. Kostarelos, Graphene materials as 2D non-viral gene transfer vector platforms, Gene Ther., 2017, 24, 123–132.

3. H. Y. Mao, S. Laurent, W. Chen, O. Akhavan, M. Imani, A. A. Ashkarran and M. Mahmoudi, Graphene: Promises, Facts, Opportunities, and Challenges in Nanomedicine, Chem. Rev., 2013, 113, 3407–3424.

4. A. F. Rodrigues, L. Newman, N. Lozano, S. P. Mukherjee, B. Fadeel, C. Bussy and K. Kostarelos, A blueprint for the synthesis and characterisation of thin graphene oxide with controlled lateral dimensions for biomedicine, 2D Mater., 2018, 5, 035020.

5. Y. Qu, F. He, C. Yu, X. Liang, D. Liang, L. Ma and Q. Zhang, Advances on graphene-based nanomaterials for biomedical applications, Mater. Sci. Eng. C, 2018, 90, 764–780.

6. E. S. Shibu, M. Hamada, N. Murase and V. Biju, Nanomaterials formulations for photothermal and photodynamic therapy of cancer, Journal Photochem. Photobiol. C Photochem. Rev., 2013, 15, 53–72.

7. N. Kundu, D. Mukherjee, T. K. Maiti and N. Sarkar, Protein-Guided Formation of Silver Nanoclusters and Their Assembly with Graphene Oxide as an Improved Bioimaging Agent with Reduced Toxicity, J. Phys. Chem. Lett, 2017, 8, 2291–2297.

8. X. Ma, H. Tao, K. Yang, L. Feng, L. Cheng, X. Shi, Y. Li, L. Guo and Z. Liu, A functionalized graphene oxide-iron oxide nanocomposite for magnetically targeted drug delivery, photothermal therapy, and magnetic resonance imaging, Nano Res., 2012, 5, 199–212.

9. J. Tian, L. Ding, Q. Wang, Y. Hu, L. Jia, J. Yu and H. Ju, Folate Receptor-Targeted and Cathepsin B-Activatable Nanoprobe for *In Situ* Therapeutic Monitoring of Photosensitive Cell Death, Anal. Chem. 2015, 87, 3841–3848.

10. Q. Liu, N. Li, M. Wang, L. Wang and X. Su, A label-free fluorescent biosensor for the detection of protein kinase activity based on gold nanoclusters / graphene oxide hybrid materials, Anal. Chim. Acta, 2018, 1013, 71–78.

11. A. M. Díez-Pascual and A. L. Díez-Vicente, Poly(propylene fumarate)/Polyethylene Glycol-Modified Graphene Oxide Nanocomposites for Tissue Engineering, ACS Appl. Mater. Interfaces, 2016, 8, 17902–17914.

12. A. Akinc and G. Battaglia, Exploiting Endocytosis for Nanomedicines, Cold Spring Harb. Perspect. Biol., 2013, 5, a016980.

13. R. Imani, F. Mohabatpour and F. Mostafavi, Graphene-based Nano-Carrier modifications for gene delivery applications, Carbon, 2018, 140, 569–591.

14. I. Canton and G. Battaglia, Endocytosis at the nanoscale, Chem. Soc. Rev., 2012, 41, 2718–2739.

15. Y. T. Fong, C.-H. Chen and J.-P. Chen, Intratumoral Delivery of Doxorubicin on Folate-Conjugated Graphene Oxide by In-Situ Forming Thermo-Sensitive Hydrogel for Breast Cancer Therapy, Nanomaterials, 2017, 7, 388.

16. S. K. Tripathi, R. Goyal, K. C. Gupta and P. Kumar, Functionalized graphene oxide mediated nucleic acid delivery, Carbon, 2013, 51, 224–235.

17. X. C. Qin, Z. Y. Guo, Z. M. Liu, W. Zhang, M. M. Wan and B. W. Yang, Folic acid-conjugated graphene oxide for cancer targeted chemo-photothermal therapy, J. Photochem. Photobiol. B Biol., 2013, 120, 156–162.

18. B. Wang, X. Su, J. Liang, L. Yang, Q. Hu, X. Shan, J. Wan and Z. Hu, Synthesis of polymer-functionalized nanoscale graphene oxide with different surface charge and its cellular uptake, biosafety and immune responses in Raw264.7 macrophages, Mater. Sci. Eng. C, 2018, 90, 514–522.

19. N. Ma, J. Liu, W. He, Z. Li, Y. Luan, Y. Song and S. Garg, Folic acid-grafted bovine serum albumin decorated graphene oxide: An efficient drug carrier for targeted cancer therapy, J. Colloid Interface Sci., 2017, 490, 598–607.

20. H. Zhu, B. Zhou, L. Chan, Y. Du and T. Chen, Transferrin-functionalized nanographene oxide for delivery of platinum complexes to enhance cancer-cell selectivity and apoptosis-inducing efficacy, Int. J. Nanomedicine, 2017, 12, 5023–5038.

21. A. E. Nel, L. Mädler, D. Velegol, T. Xia, E. M. V Hoek, P. Somasundaran, F. Klaessig, V. Castranova and M. Thompson, Understanding biophysicochemical interactions at the nano-bio interface., Nat. Mater., 2009, 8, 543–57.

22. J. Rejman, V. Oberle, I. S. Zuhorn and D. Hoekstra, Size-dependent internalization of particles via the pathways of clathrin- and caveolae-mediated endocytosis, Biochem. J., 2004, 377, 159–169.

23. J. Huang, C. Zong, H. Shen, M. Liu, B. Chen, B. Ren and Z. Zhang, Mechanism of Cellular Uptake of Graphene Oxide Studied by Surface-Enhanced Raman Spectroscopy, Small, 2012, 8, 2577–2584.

24. Q. Mu, G. Su, L. Li, B. O. Gilbertson, L. H. Yu, Q. Zhang, Y.-P. Sun and B. Yan, Size-Dependent Cell Uptake of Protein-Coated Graphene Oxide Nanosheets, ACS Appl, Mater. Interfaces, 2012, 4, 2259–2266.

25. J. Linares, M. C. Matesanz, M. Vila, M. J. Feito, G. Goncalves, M. Vallet-Regi, P. A. A. P. Marques, M. T. Portoles, Endocytic Mechanisms of Graphene Oxide Nanosheets in Osteoblasts, Hepatocytes and Macrophages, ACS Appl. Mater. Interfaces, 2014, 6, 13697–13706.

26. V. Francia, C. Reker-Smit, G. Boel, A. Salvati, Limits and challenges in using transport inhibitors to characterize how nano-sized drug carriers enter cells, Nanomedicine (Lond.), 2019, 14 (12), 1533 – 1549.

27. S. Vranic, A. F. Rodrigues, M. Buggio, L. Newman, M. R. H. White, D. G. Spiller, C. Bussy and K. Kostarelos, Live Imaging of Label-Free Graphene Oxide Reveals Critical Factors Causing Oxidative Stress-Mediated Cellular Responses, ACS Nano, 2018, 12, 1373–1389.

28. L. S. Franqui, M. A. De Farias, R. V. Portugal, C. A. R. Costa, R. R. Domingues, A. G. Souza Filho, V. R. Coluci, A. F. P. Leme and D. S. T. Martinez, Interaction of graphene oxide with cell culture medium: Evaluating the fetal bovine serum protein corona formation towards in vitro nanotoxicity assessment and nanobiointeractions, Mater. Sci. Eng. C, 2019, 100, 363–377.

29. C. Bussy and K. Kostarelos, Culture Media Critically Influence Graphene Oxide Effects on Plasma Membranes, Chem, 2017, 2, 322–323.

30. T. L. Moore, L. Rodriguez-Lorenzo, V. Hirsch, S. Balog, D. Urban, C. Jud, B. Rothen-Rutishauser, M. Lattuada and A. Petri-Fink, Nanoparticle colloidal stability in cell culture media and impact on cellular interactions, Chem. Soc. Rev., 2015, 44, 6287–6305.

31. G. Duan, S. Kang, X. Tian, J. A. Garate, L. Zhao, C. Ge, R. Zhou, Protein corona mitigates the cytotoxicity of graphene oxide by reducing its physical interaction with cell membrane, Nanoscale, 2015, 7, 15214–15224.

32. C. Hermans and A. Bernard, Lung Epithelium–specific Proteins, Characteristics and Potential Applications as Markers, Am. J. Respir. Crit. Care Med. 1999, 159, 646–678.

33. K.-J. Kim and A. B. Malik, Protein transport across the lung epithelial barrier. Am. J. Physiol Lung Cell Mol. Physiol, 2003, 284, L247–L259.

34. H. J. Issaq, Z. Xiao and T. D. Veenstra, Serum and Plasma Proteomics, Chem. Rev., 2007, 107, 3601–3620.

35. C. Jin, F. Wang, Y. Tang and X. Zhang, J. Wang, Y. Yang, Distribution of Graphene Oxide and TiO_2_-Graphene Oxide Composite in A549 Cells, Biol. Trace Elem. Res., 2014, 159, 393–398.

36. R. G. Mendes, B. Koch, A. Bachmatiuk, X. Ma, S. Sanchez, C. Damm, O. G. Schmidt, T. Gemming, J. Eckert and M. H. Rümmeli, A size dependent evaluation of the cytotoxicity and uptake of nanographene oxide, J. Mater. Chem. B, 2015, 3, 2522–2529.

37. Y. Chang, S.-T. Yang, J.-H. Liu, E. Dong, Y. Wang, A. Cao, Y. Liu and H. Wang, In vitro toxicity evaluation of graphene oxide on A549 cells, Toxicol. Lett., 2011, 200, 201–210.

38. R. G. Mendes, A. Mandarino, B. Koch, A. K. Meyer, A. Bachmatiuk, C. Hirsch, T. Gemming, O. G. Schmidt, Z. Liu and M. H. Rümmeli, Size and time dependent internalization of label-free nano-graphene oxide in human macrophages, Nano Res., 2017, 10, 1980–1995.

39. A. Lesniak, F. Fenaroli, M. P. Monopoli, C. Åberg, K. A. Dawson and A. Salvati, Effects of the presence or absence of a protein corona on silica nanoparticle uptake and impact on cells, ACS Nano, 2012, 6, 5845–5857.

40. M. P. Monopoli, C. Åberg, A. Salvati and K. A. Dawson, Biomolecular coronas provide the biological identity of nanosized materials, Nat. Nanotechnol., 2012, 7, 779–786.

41. M. Lotya, A. Rakovich, J. F. Donegan and J. N. Coleman, Measuring the lateral size of liquid-exfoliated nanosheets with dynamic light scattering, Nanotechnology, 2013, 24, 265703.

42. R. D. Santo, L. Digiacomo, S. Palchetti, V. Palmieri, G. Perini, D. Pozzi, M. Papi and G. Caracciolo, Microfluidic manufacturing of surface-functionalized graphene oxide nanoflakes for gene delivery, Nanoscale, 2019, 11, 2733–2741.

43. E. Bidram, A. Sulistio, A. Amini, Q. Fu, G. G. Qiao, A. Stewart and D. E. Dunstan, Fractionation of graphene oxide single nano-sheets in water-glycerol solutions using gradient centrifugation, Carbon, 2016, 103, 363–371.

44. S. Bhattacharjee, DLS and zeta potential - What they are and what they are not?, J. Control. Release, 2016, 235, 337–351.

45. M. S. Ehrenberg, A. E. Friedman, J. N. Finkelstein, Güter Oberdörster and J. L. Mcgrath, The influence of protein adsorption on nanoparticle association with cultured endothelial cells, Biomaterials, 2009, 30, 603–610.

46. L. S. Franqui, M. A. De Farias, R. V. Portugal, C. A. R. Costa, R. R. Domingues, A. G. Souza Filho, V. R. Coluci, A. F. P. Leme and D. S. T. Martinez, Interaction of graphene oxide with cell culture medium: Evaluating the fetal bovine serum protein corona formation towards in vitro nanotoxicity assessment and nanobiointeractions, Mater. Sci. Eng. C, 2019, 100, 363–377.

47. D. Mahl, C. Greulich, W. Meyer-Zaika, M. Köller and M. Epple, Gold nanoparticles: dispersibility in biological media and cell-biological effect, J. Mater. Chem., 2010, 20, 6176–6181.

48. H. Hillaireau and P. Couvreur, Nanocarriers’ entry into the cell: relevance to drug delivery, Cell. Mol. Life Sci., 2009, 66, 2873–2896.

49. I. M. Martinez Paino, F. Santos and V. Zucolotto, Biocompatibility and toxicology effects of graphene oxide in cancer, normal, and primary immune cells, J. Biomed. Mater. Res. Part A, 2017, 105A, 728–736.

50. M. I. Setyawati, C. Y. Tay, D. Docter, R. H. Stauber and D. T. Leong, Understanding and exploiting nanoparticles’ intimacy with the blood vessel and blood, Chem. Soc. Rev., 2015, 44, 8174–8199.

51. J. P. Lim and P. A. Gleeson, Macropinocytosis: an endocytic pathway for internalising large gulps, Immunol. Cell Biol., 2011, 89, 836–843.

52. Z. Kadlecova, S. J. Spielman, D. Loerke, A. Mohanakrishnan, D. K. Reed and S. L. Schmid, Regulation of clathrin-mediated endocytosis by hierarchical allosteric activation of AP2, J. Cell Biol., 2017, 216, 167–179.

53. M. Kaksonen and A. Roux, Mechanisms of clathrin-mediated endocytosis, Nat. Rev. Mol. Cell Biol., 2018, 19, 313–326.

54. E. Cocucci, F. Aguet, S. Boulant and T. Kirchhausen, The first five seconds in the life of a clathrin-coated pit, Cell, 2012, 150, 495–507.

55. G. J. Doherty and H. T. McMahon, Mechanisms of Endocytosis, Annu. Rev. Biochem., 2009, 78, 857–902.

56. R. G. Parton and K. Simons, The multiple faces of caveolae, Nat. Rev. Mol. Cell Biol., 2007, 8, 185–194.

57. D. Vercauteren, R. E. Vandenbroucke, A. T. Jones, J. Rejman, J. Demeester, S. C. De Smedt, N. N. Sanders and K. Braeckmans, The Use of Inhibitors to Study Endocytic Pathways of Gene Carriers: Optimization and Pitfalls, Mol. Ther., 2010, 18, 561–569.

58. D. A. Kuhn, D. Vanhecke, B. Michen, F. Blank, P. Gehr, A. Petri-Fink and B. Rothen-Rutishauser, Different endocytotic uptake mechanisms for nanoparticles in epithelial cells and macrophages, Beilstein J. Nanotechnol., 2014, 5, 1625–1636.

59. K. L. Douglas, C. A. Piccirillo and M. Tabrizian, Cell line-dependent internalization pathways and intracellular trafficking determine transfection efficiency of nanoparticle vectors, 2008, 68, 676–687.

60. E. Macia, M. Ehrlich, R. Massol, E. Boucrot, C. Brunner and T. Kirchhausen, Dynasore, a Cell-Permeable Inhibitor of Dynamin, Dev. Cell, 2006, 10, 839–850.

61. C.-H. Wu, Y.-P. Chen, S.-H. Wu, Y. Hung, C.-Y. Mou and R. P. Cheng, Enhanced Non-Endocytotic Uptake of Mesoporous Silica Nanoparticles by Shortening the Peptide Transporter Arginine Side Chain, ACS Appl. Mater. Interfaces, 2013, 5, 12244–12248.

62. A. Verma, O. Uzun, Y. Hu, Y. Hu, H. Han, N. Watson, S. Chen, D. J. Irvine and F. Stellacci, Surface-structure-regulated cell-membrane penetration by monolayer-protected nanoparticles, Nat. Mater., 2008, 7, 588–595.

63. W. Bao, J. Wang, Q. Wang, D. O. Hare and Y. Wan, Layered Double Hydroxide Nanotransporter for Molecule Delivery to Intact Plant Cells, Sci. Rep., 2016, 6, 26738.

64. Q. Mu, N. S. Hondow, Ł. Krzemi, A. P. Brown, L. J. C. Jeuken and M. N. Routledge, Mechanism of cellular uptake of genotoxic silica nanoparticles, PART. FIBRE. TOXICOL., 2012, 9, 1–11.

65. S. C. Silverstein, R. M. Steinman and Z. A. Cohn, Endocytosis, Ann. Rev. Biochem. 1977, 46, 669–722.

66. R. M. Steinman, J. M. Silver and Z. A. Cohn, PINOCYTOSIS IN FIBROBLASTS Quantitative Studies In Vitro, J. Cell Biol., 1974, 63, 949–969.

67. S. Subramanya, C. F. Hardin, D. Steverding and K. Menda-Wilmot, Glycosylphosphatidylinositol-specific phospholipase C regulates transferrin endocytosis in the African trypanosome, Biochem J., 2009, 417, 685–694.

68. H. Zhang, C. Peng, J. Yang, M. Lv, R. Liu, D. He, C. Fan and Q. Huang, Uniform Ultrasmall Graphene Oxide Nanosheets with Low Cytotoxicity and High Cellular Uptake, ACS Appl. Mater. Interfaces, 2013, 5, 1761–1767.

69. D. Dutta and J. G. Donaldson, Search for inhibitors of endocytosis: Intended specificity and unintended consequences., Cell. Logist., 2012, 2, 203–208.

70. L. M. Fujimoto, R. Roth, J. E. Heuser and S. L. Schmid, Actin Assembly Plays a Variable, but not Obligatory Role in Receptor-Mediated Endocytosis, Traffic, 2000, 1, 161–171.

71. S. Ka, B. Amstutz, M. Gastaldelli, N. Wolfrum, K. Boucke, M. Havenga, F. Digennaro, N. Liska, S. Hemmi and U. F. Greber, Macropinocytotic Uptake and Infection of Human Epithelial Cells with Species B2 Adenovirus Type 35, J Virol, 2010, 84, 5336–5350.

72. B. Alberts, A. Johnson, J. Lewis, M. Raff, K. Roberts, P. Walter, Molecular Biology of the Cell, 6th edn, 2014.

73. A. I. Ivanov, Pharmacological inhibitors of exocytosis and endocytosis: Novel bullets for old targets, Methods Mol. Biol., 2014, 1174, Chapter 1.

74. T. dos Santos, J. Varela, I. Lynch, A. Salvati and K. A. Dawson, Effects of Transport Inhibitors on the Cellular Uptake of Carboxylated Polystyrene Nanoparticles in Different cell lines, PLoS One, 2011, 6, e24438.

75. T. Xia, M. Kovochich, M. Liong, J. I. Zink and A. E. Nel, Cationic Polystyrene Nanosphere Toxicity Depends on Cell-specific Endocytic and Mitochondrial Injury Pathways, ACS Nano, 2008, 2, 85–96.

76. R. K. Srivastava, C. Li, J. Khan, N. S. Banerjee, L. T. Chow and M. Athar, Combined mTORC1/mTORC2 inhibition blocks growth and induces catastrophic macropinocytosis in cancer cells, Proc. Natl. Acad. Sci., 2019, 119 (49), 24583–24592.

77. S. Shibutani, H. Okazaki and H. Iwata, Dynamin-dependent amino acid endocytosis activates mechanistic target of rapamycin complex 1 (mTORC1), J. Biol. Chem., 2017, 292 (44), 18052–18061.

78. M. Jin, Z. Liu, W. Zhang, H. Dong, F. Zhou, J. Yu, X. Wang and Z. Guo, Mitochondrial-Targeted Polyethylenimine Functionalized Graphene Oxide Nanocarrier and its Anti-Tumor Effect on Human Lung Carcinoma Cells, Nano, 2015, 10, 1550121.

79. D. A. Jasim, N. Lozano, K. Kostarelos, Synthesis of few-layered, high-purity graphene oxide sheets from different graphite sources for biology, 2D Mater., 2016, 3, 014006.

80. M. I. Love, W. Huber and S. Anders, Moderated estimation of fold change and dispersion for RNA-seq data with DESeq2, Genome Biol.,2014, 15, 550.

81. R Core Team, R: A language and environment for statistical computing. R Foundation for Statistical Computing, Vienna, Austria, 2018, URL https://www.R-project.org/

82. W. Jawaid, enrichR: Provides an R Interface to ‘Enrichr’, 2019, R package version 2.1, https://CRAN.R-project.org/package=enrichR

83. R. Kolde, pheatmap: Pretty Heatmaps, 2019, R package version 1.0.12, https://CRAN.R-project.org/package=pheatmap

